# Leptin reduces pathology and increases adult neurogenesis in a transgenic mouse model of Alzheimer’s disease

**DOI:** 10.1101/567636

**Authors:** Michele Longoni Calió, Geisa Nogueira Salles, Darci Souza Marinho, Amanda Fávero Mosini, Fernando Henrique Massinhani, Gui Mi Ko, Marimélia A. Porcionatto

**Author notes:** Corresponding Authors: Marimélia A. Porcionatto, Michele Longoni Calió, Rua Pedro de Toledo, 669 – 3^°^ andar, CEP 04049-032, São Paulo - SP – Brazil.

## Abstract

Alzheimer’s disease (AD) is the most common dementia worldwide, and is characterized by the presence, in the brain tissue, of extracellular senile plaques formed by amyloid beta (Aβ) peptide and intracellular neurofibrillary tangles of hyperphosphorylated Tau protein. These changes lead to progressive neuronal degeneration and dysfunction, resulting in severe brain atrophy and cognitive deficits. With the discovery that neurogenesis persists in the adult mammalian brain, including brain regions affected by AD, studies of the use of neural stem cells for treatment of neurodegenerative diseases in order to repair and/or prevent neuronal cell loss have increased. Here we show that leptin increases neurogenesis in the dentate gyrus of adult mice as well as in the subventricular zone both in wild type and AD transgenic mouse model. Chronic administration of leptin to young mice increased neural stem cell proliferation with significant effects on differentiation and survival of newborn cells. Expression of the long form of leptin receptor, LepRb, was detected in the neurogenic niches by reverse transcription-PCR and immunohistochemistry. Moreover, leptin modulated astrogliosis and the formation of senile plaques. Additionally, leptin led to attenuation of Aβ-induced neurodegeneration and superoxide anion production as revealed by Fluoro-Jade B and dihydroethidium (DHE) staining. Our study contributes to the understanding of the effects of leptin in the brain that may lead to the development of new therapies to treat Alzheimer’s disease.

## INTRODUCTION

Described over a century ago, Alzheimer’s disease (AD) is the most common progressive dementia in late adult life (Dai et al., 2017; Gallaway et al, 2017) and still is not fully understood. The pathology of AD is mainly characterized by multiple etiological and biochemical aberrations including the formation of senile plaques by aggregates of amyloid beta (Aβ) peptides, neurofibrillary tangles (NFTs) of hyperphosphorylated Tau protein and oxidative stress, that are linked to abnormal neuronal function and progressive degeneration and neuronal loss (Zhang, 2011; Armstrong, 2011; Lauritzen et al., 2012; Reitz and Mayeux, 2014; Jeong, 2017; Sanabria-Castro et al., 2017; Feitosa, 2018).

It is well recognized that the proliferative activity of neural stem cells (NSC) markedly decreases during aging and is dysregulated in neurodegenerative diseases. The discovery that quiescent precursor cells can be activated even in the aged or damaged brain has contributed to the emergence of the hypothesis that neurodegenerative diseases such as AD could be treated by stimulating the neurogenic potential, and the derived progeny might be re-directed to areas of degeneration, where they would be involved in cell replacement (for reviews, see Gage, 2002; Hormoz, 2013; Matarredona et al., 2018). Consequently, several studies have been conducted to investigate adult neurogenesis in AD (Taniuchi et al., 2007; Naumann et al., 2010; Kanemoto et al., 2014; Hamilton et al., 2015; Winner and Winkler, 2015; Tai et al., 2017).

Clinical evidence indicates that diet and lifestyle are major risk factors for developing AD and disturbances in metabolism are also linked to the development of the pathology (Dosunmu et al., 2007; Yin et al., 2016). The hormone leptin is known to regulate food intake and energy balance, being also involved in many other processes such as immune response, stress, and neurodevelopment (Folch et al., 2012; Bellefontaine et al., 2014; Meyer et al., 2014). Indeed, lower leptin levels are associated with an increased risk of developing AD, as well as aberrant leptin function (Lieb et al., 2009; Baranowska-Bik et al., 2015). Within the past decade, researchers showed a role for leptin in neuroprotection and hippocampal neurogenesis. In animal models, leptin administration facilitates learning and alter synaptic plasticity (Moult et al., 2010; Luo et al., 2015; Ghasemi et al., 2016). Additionally, this adipokine attenuates Tau hyperphosphorylation in neuronal cells and protects against Aβ-induced neurotoxicity in cellular models (Garza et al. 2008, Doherty et al., 2013; Liu et al., 2015). Thus, there is growing evidence that leptin-based therapies may be beneficial in AD.

We used the double transgenic APPswe/PS1dE9 (2xTgAD) mouse model for investigating the action of leptin in two neurogenic niches. The 2xTgAD mouse lineage reproduces a subset of neuropathological, biochemical and behavioral changes identified in patients with AD, such as massive Aβ accumulation and plaque formation, vascular deficits and memory impairment (Holcomb et al., 1998; Jankowsky et al., 2002).

To further characterize the association between leptin and neurogenesis in AD, we assessed the presence of the long form of leptin receptor (LepRb) in 2xTgAD and wild-type (WT) leptin-treated mice at different ages and analyzed the co-localization of LepRb with the markers of proliferation and neural differentiation in two neurogenic niches, the subventricular zone (SVZ) lining the lateral ventricles and the subgranular zone (SGZ) of the hippocampal dentate gyrus. We also evaluated the production of reactive oxygen species (ROS), the formation of Aβ plaques and neurodegeneration in leptin-treated and untreated mice. Interestingly, we observed that leptin was able to increase proliferation and reduce senile plaque formation, neuronal death, and astrogliosis even in the aged 2xTgAD animals. Moreover, the significant decrease in superoxide anion (O_2_^−^) production observed after treatment indicates that leptin also has antioxidant action in AD. The overexpression of LepRb suggests its involvement in the observed improvement.

As far as we know, our findings are the first to demonstrate that leptin plays a role in extra-hippocampal neurogenesis, in regions involved in neural cell replacement, regardless of age or disease stage. Collectively, we show that leptin participates in several events that decrease the progression of the disease and exert multiple functions besides neurogenesis, by decreasing senile plaque formation and oxidative stress. We hypothesized that strategies targeted at stimulating this process could have significant therapeutic value, suggesting that leptin may be used as a strategy for the treatment of AD and probably other neurodegenerative diseases.

## RESULTS

### Leptin Changes Gene Expression in the Hippocampus

We treated WT and 2xTgAD mice with leptin for 7 consecutive days (See Supplemental Data Figure S1 A). Body weight of the mice was measured before and throughout the 7 days of leptin treatment and the data of each group recorded daily are shown in Spplemental Data Figure S01B and the percentage of body weight loss are show in Figure S 01D. The treatment with leptin had no effect on weight loss of animals.

We observed that there is a tendency in the decrease of serum leptin concentration in 2xTgAD animals, although there was no significant difference. On the other hand, we did not detect a significant enhancement in leptin levels in serum samples in 2xTgAD mice after leptin treatment (See Suplemental Data Figure S01C). Both results may be due to leptin half-life in plasma, which is about 40 min. These results also suggest a very restricted and subtle target (in brain), preventing possible peripheral side effects.

After this, we investigated the expression profile of genes involved in neurogenesis in the hippocampus: glial fibrillary acid protein (GFAP - neural stem cell and astrocyte marker); Nestin (amplifying and neural stem cell marker); proliferating cell nuclear antigen (PCNA) and Ki67 (proliferation and cell-cycle markers); doublecortin (DCX - neuroblast marker); βIII-tubulin, microtubule associated protein2a (Map2a) and neuronal specific nuclear protein (NeuN - mature neuron markers).

We observed that treatment with leptin increased the expression of Nestin by young WT, and young and aged 2xTgAD mice (Figure 1B). Expression of DCX was increased in all treated groups (Figure 1E) whereas we observed upregulation of PCNA in young animals of both strains (Figure 1C). GFAP expression was downregulated in leptin-treated aged 2xTgAD animals, but not in the other groups (Figure 1A). Finally, no difference was observed in the expression of tubulin βIII, Map2a and NeuN with the use of leptin, but a decrease in the expression of these genes was found in aged 2xTgAD mice in comparison to the WT or young animals (Figure 1F-H).

**Figure 1.**
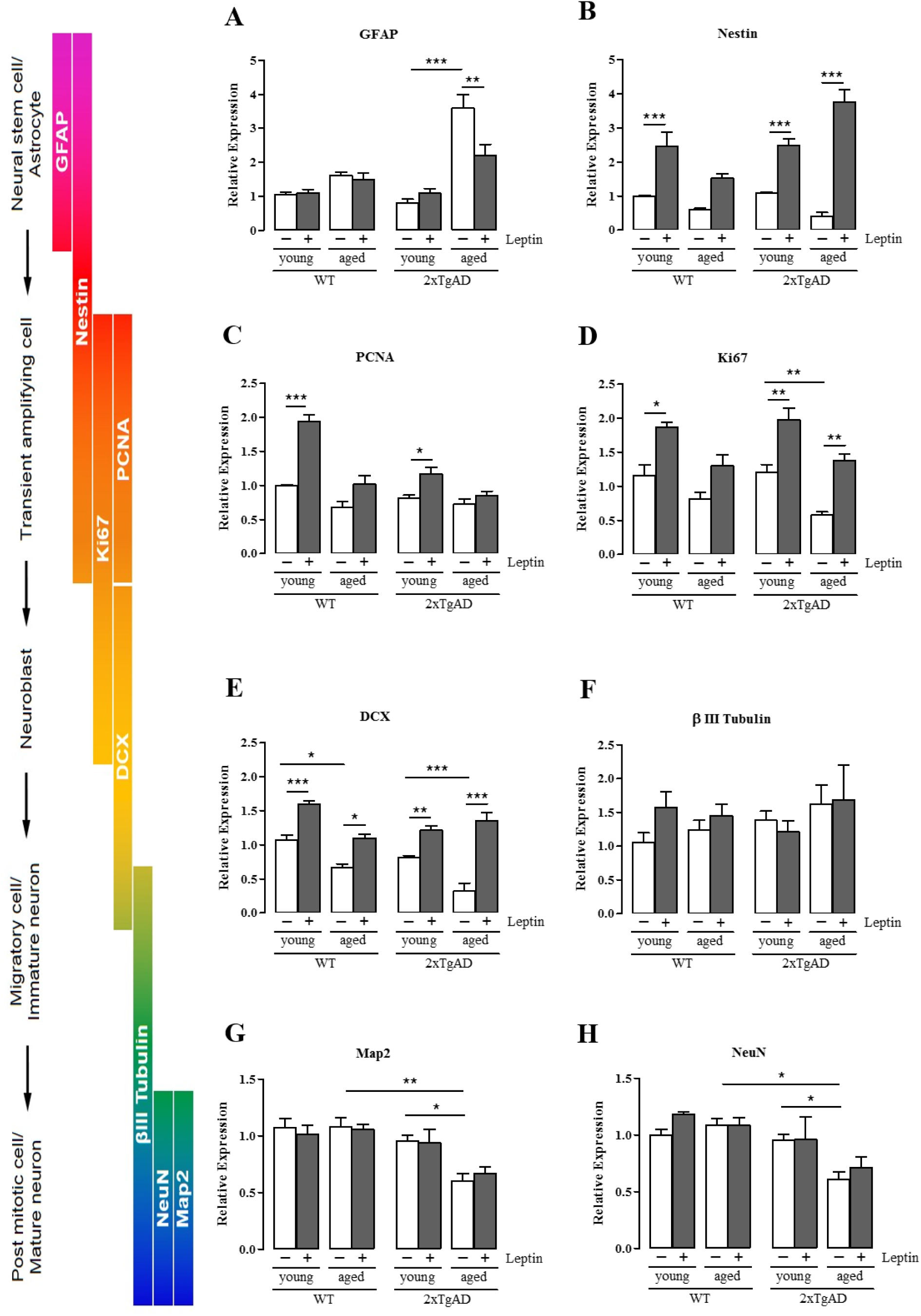
Gene expression analysis by qPCR of proliferative and neural markers in the hippocampus of mice treated or not with leptin in different ages. The animals were injected i.p. with vehicle (PBS) or leptin (1mg/kg) daily for 7 days. Leptin treatment increased the expression of most genes analyzed. Notice the difference in the expression of GFAP **(A)**, Ki67 **(D)** and DCX **(E)** between young and aged mice. The data were normalized using four endogenous controls and are relative to young WT PBS control group; *n* = 6 per group. One-way ANOVA, data are shown as mean ± SEM, represented by error bars; \**p* < 0.05, \*\**p* < 0.01 \*\*\**p* < 0.001.

### Leptin increases proliferation of neuronal progenitors

In adult brain, neurogenesis occurs in the SGZ of the hippocampal dentate gyrus and in the SVZ of the lateral ventricle. While migration of the new generated cells in the SGZ is restricted to the hippocampus, where they participate in learning and memory formation, cells from the SVZ are destined to renew interneurons of the olfactory bulb (Kriegstein and Alvarez-Buylla, 2009). SVZ-migrating cells are thought to participate in the attempt to regenerate an injured area (Conover and Notti, 2008; Clelland et al., 2009; Deng et al., 2009; Garthe et al., 2009; Mundim et al, 2018). But with increasing age, the neurogenesis process declines, impacting its physiological role in neuronal replacement and consequently its ability to participate in the repair mechanisms triggered by injury or disease (Enwere et al., 2004).

Due to the fact that the hippocampus is affected in AD, most studies about neuroregeneration examine the involvement of SGZ-born neuroblasts in brain repair, and neurogenesis at the SVZ has not been investigated. Therefore, in the present study, we asked whether leptin regulates adult neurogenesis in the SVZ and in the hippocampus. To address this question, we labelled newborn cells by administrating 5-bromo-2’deoxyuridine (BrdU) in the last two days of treatment with leptin, and quantified the number of BrdU^+^ cells in the SGZ of hippocampus and in the SVZ of treated and untreated, young and aged, WT and 2xTgAD mice.

In the hippocampus, BrdU^+^ cells usually are distributed in the inner layer of the granular cell layer of the dentate gyrus, while in SVZ, neural stem cells can be found in several areas, although we have focused on the dorsolateral region (Figure 2A). We found significantly higher number of BrdU^+^ cells in the SVZ of all leptin-treated groups (Figure 2B). Analysis of the SGZ revealed that all groups, except the young WT mice showed increased number of BrdU^+^ cells following treatment with leptin (Figure 2C).

**Figure 2.**
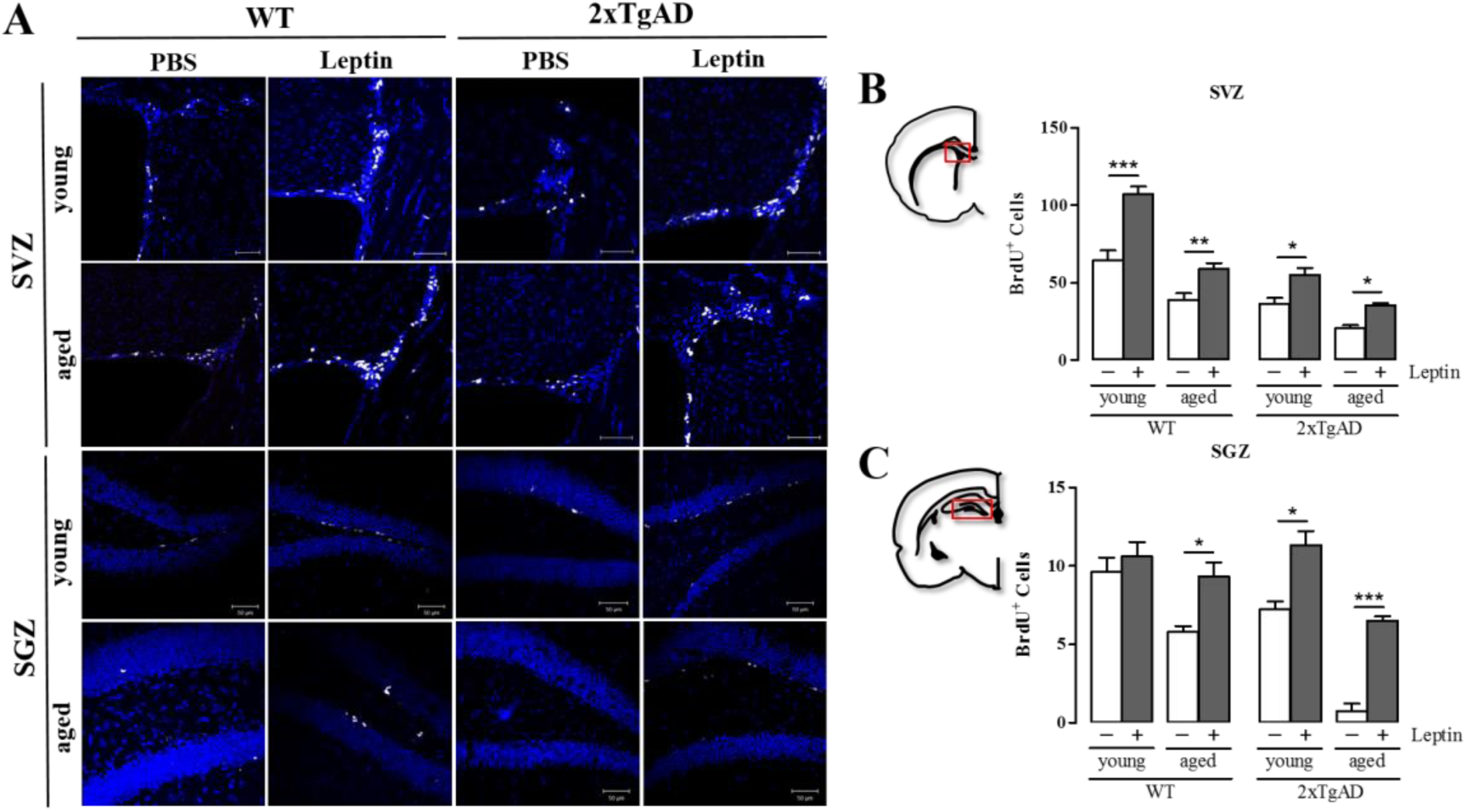
*In vivo* leptin administration increases proliferation in the SVZ and SGZ neurogenic niches. BrdU labeling was used to evaluate proliferation in both neurogenic niches. The animals were injected i.p. with vehicle (PBS) or leptin (1mg/kg) daily for 7 days. BrdU^+^ cells were counted in the SVZ of the lateral ventricle and in the SGZ of the dentate gyrus of the hippocampus. **(A)** Representative images showing BrdU^+^ cells in the SVZ and SGZ from control (PBS) and treated (leptin) WT and 2xTgAD mice of different ages. **(C)** Regions used for the analysis and quantification of BrdU^+^ cells showing increased proliferation in the treated groups in the SVZ **(B)** and SGZ **(C)**. Scale bar = 50µm. Data are expressed as mean ± SEM, *n* = 5 animals per group. \**p* < 0.05; \*\**p* < 0.01; \*\*\**p* < 0.001.

### Leptin decreases astrogliosis in the hippocampus and increases neurogenesis in the SGZ and SVZ

We observed that leptin treatment led to a decrease in the number of GFAP^+^ cells in the hippocampus of aged 2xTgAD animals (Figure 3A, C), suggesting that leptin was able to reduce astrogliosis. The number of GFAP-labeled cells in the SVZ was not significantly altered by leptin treatment in WT and transgenic mice (Figure 3A, B).

**Figure 3.**
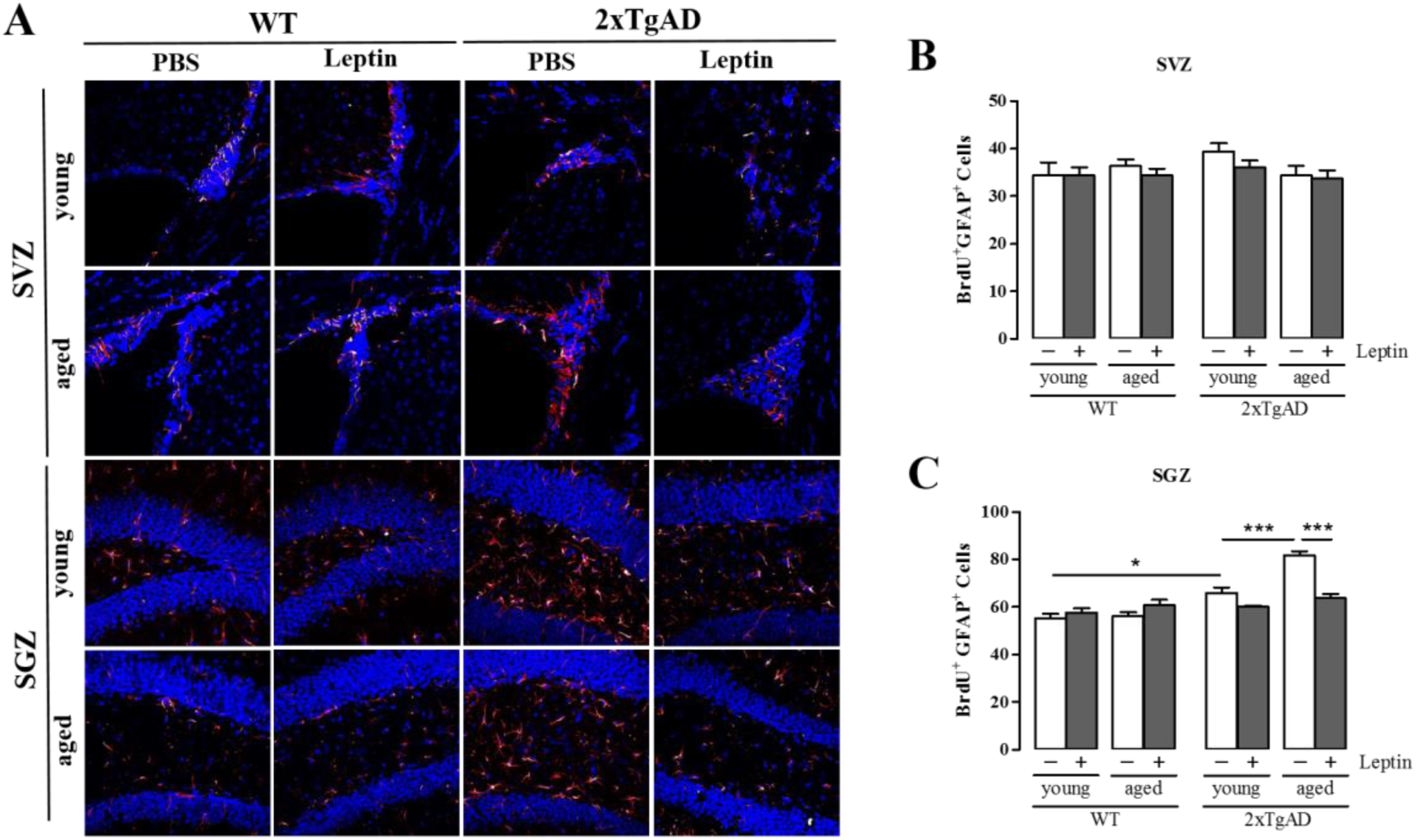
Leptin administration decreases astrogliosis in the hippocampus. GFAP labeling was used to evaluate proliferation of neural stem cell and astrocytes in both neurogenic niches. Double positive BrdU^+^/GFAP^+^ cells were counted in the SVZ of the lateral ventricle and in the hippocampus. **(A)** Representative images showing BrdU^+^ (white) and GFAP^+^ (red) cells in SVZ and SGZ from control (PBS) and treated (leptin) WT and 2xTgAD mice of different ages. **(B)** Quantification of BrdU^+^/GFAP^+^ cells shows no difference in the proliferation of neural stem cells in the SVZ when mice received leptin. **(C)** Quantification of BrdU^+^/GFAP^+^ cells show decreased proliferation of astrocyte in the hippocampus of the leptin treated group. Scale bar = 50µm. Data are expressed as mean ± SEM, *n* = 6 animals per group. \**p* < 0.05; \*\**p* < 0.01; \*\*\**p* < 0.001.

Our results show that after 7 days of leptin treatment there was a significant increase in the number of DCX^+^ cells in all groups of animals (Figure 4). Higher numbers of DCX^+^ cells were observed in both regions of interest including the dorsolateral area in SVZ and SGZ in the hippocampal dentate gyrus (Figure 4). Curiously, the presence of neuroblasts is more evident in the SVZ, in all groups analyzed, when compared to the hippocampus, where the action of leptin was more significant in the aged 2xTgAD (Figure 4C). Thus, the increments in the number of neuroblasts were approximately 1.5 fold in the SVZ and 30 fold in the aged 2xTgAD group in the hippocampus.

**Figure 4.**
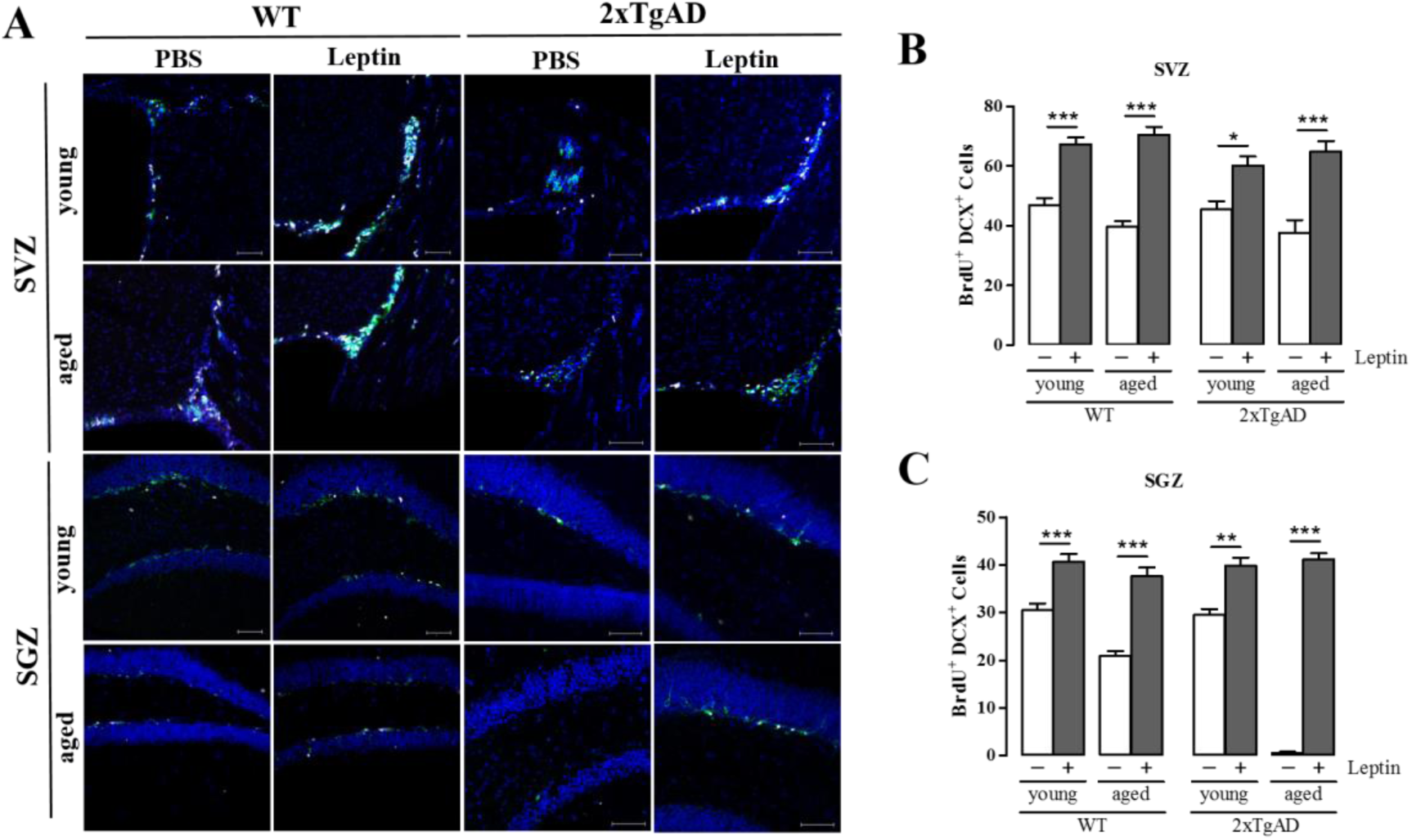
Leptin administration increases the generation of neuroblasts in the SVZ and hippocampal SGZ. DCX labeling was used to evaluate the production of neuroblasts by neural stem cells in the two neurogenic niches. Double positive BrdU^+^/DCX^+^ cells were counted in the SVZ of the lateral ventricle and in the SGZ of dentate gyrus of the hippocampus. **(A)** Representative images showing BrdU^+^ (white) and DCX^+^ (green) cells in the SVZ and SGZ from control (PBS) and treated (leptin) WT and 2xTgAD mice of different ages. **(B)** Quantification of BrdU^+^/DCX^+^ cells shows increased generation of neuroblasts in the treated groups in the SVZ. **(C)** Quantification of BrdU^+^/DCX^+^ cells show increased generation of neuroblasts in the treated groups in SGZ. Scale bar = 50µm. Data are expressed as mean ± SEM, *n* = 6 animals per group. \**p* < 0.05; \*\**p* < 0.01; \*\*\**p* < 0.001.

### Leptin treatment upregulates the expression of LepRb in adult neural stem/progenitor cells

We next sought to elucidate the mechanisms involved in transducing the effects of leptin in neurogenesis and differentiation. It has been described in the literature that LepRb is expressed in the dentate gyrus (Scott et al., 2009), and to determine if leptin treatment would affect LepRb expression, we used qPCR and immunocytochemistry.

qPCR analysis using RNA isolated from hippocampal tissue of untreated and treated groups of both ages and lineages showed that the expression level of LepRb was significantly higher in all treated groups compared to control (Figure 5A). Interestingly, there was no significant change in LepRb expression as a function of age in either WT or 2xTgAD group (Figure 5A).

**Figure 5.**
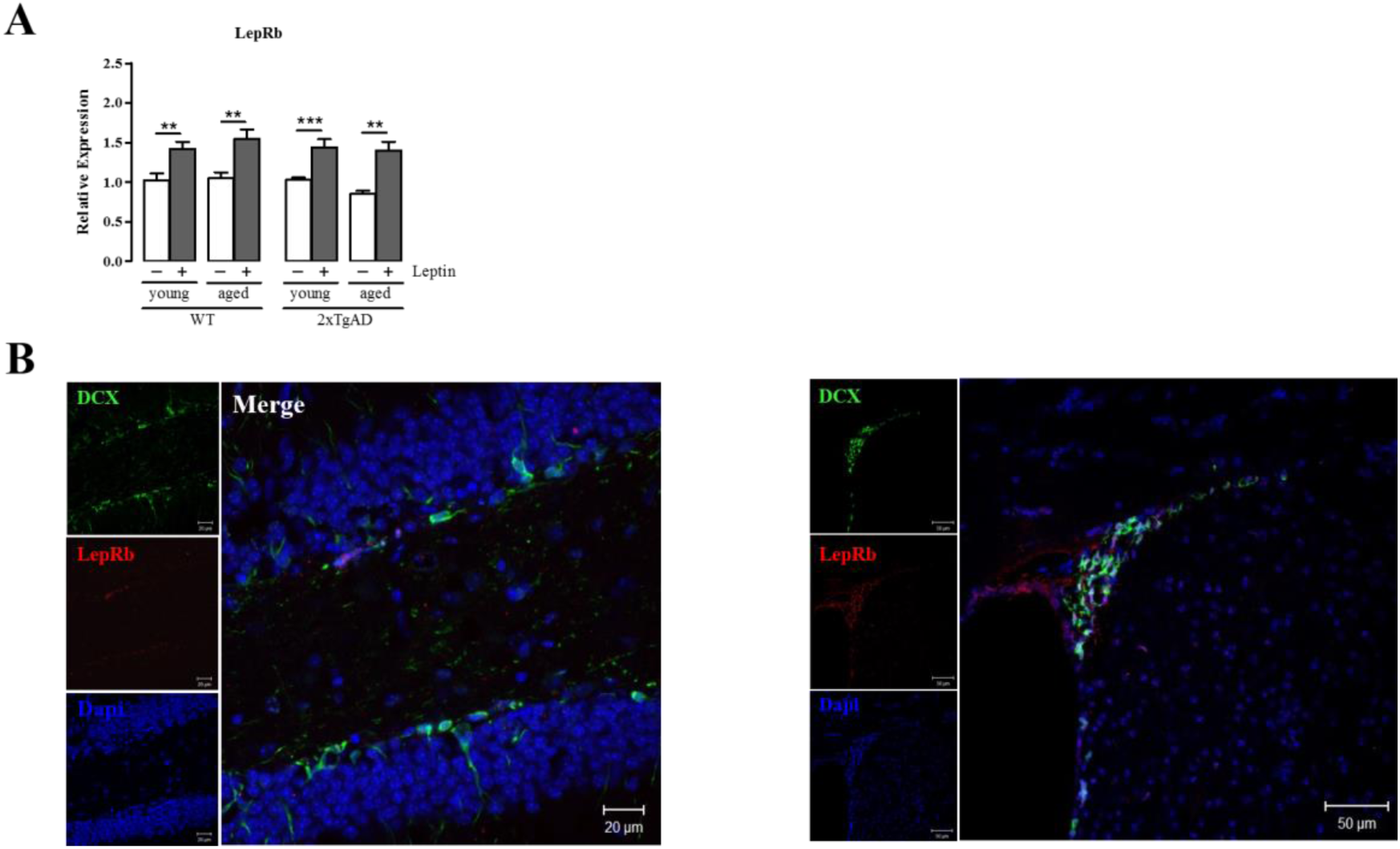
Neural stem cells and neuroblasts express LepRb. **(A)** Gene expression analysis by qPCR for LepR in the hippocampus. Leptin treatment increased the expression of the receptor on all treated groups. **(B)** Representative images show double stainning LepR^+^/DCX^+^ cells in SGZ of hippocampus. Scale bar = 20μm. **(C)** Representative images show double stainning LepR^+^/DCX^+^ cells in SVZ. Scale bar = 50μm.

To investigate which type of neural progenitor was expressing leptin receptor, the presence of LepRb on hippocampus and SVZ was further confirmed by immunohistochemical staining with antibodies against LepRb and DCX regardless of lineage or the age of animals (Figure 5B and 5C). The immunoreactivity for LepRb was observed in a large number of DCX^+^ cells. No immunostaining was seen when primary antibody was omitted.

### Leptin restores Sod2 and GPx1 expression and leads to decreased superoxide production in 2xTgAD mice

It is known that oxidative stress can initiate the process of apoptosis and that abnormalities in the regulation and expression of antioxidant enzymes may have a role in mechanisms of central nervous system (CNS) aging and neurodegeneration. In fact, neurons within the CNS are at particular risk from damage caused by free radicals because of high rates of oxygen consumption and high contents of polyunsaturated fatty acids (Zemlan et al., 1989; Uysal et al., 2012; Radi et al., 2014;). We used Custom TaqMan Array Fast Plates to Oxidative stress and Antioxidant defense (See Supplenetal Data Table S1) to investigate whether neurodegeneration and leptin treatment could affect the gene expression pattern of antioxidant enzymes. The number of upregulated or downregulated genes for each group are listed in Table 1. Most genes related to antioxidant enzymes that could have their relative expression (RE) quantified, did not show significant change in the treated groups (Table 2), except for superoxide dismutase-2 (*Sod2*) and Glutathione peroxidase-1 (*GPx1*) in 2xTgAD treated animals (Figure 6A, B). Interestingly, even in the younger DA models, the treatment with leptin induced an increase on gene expression of these two genes.

**TABLE 1:**
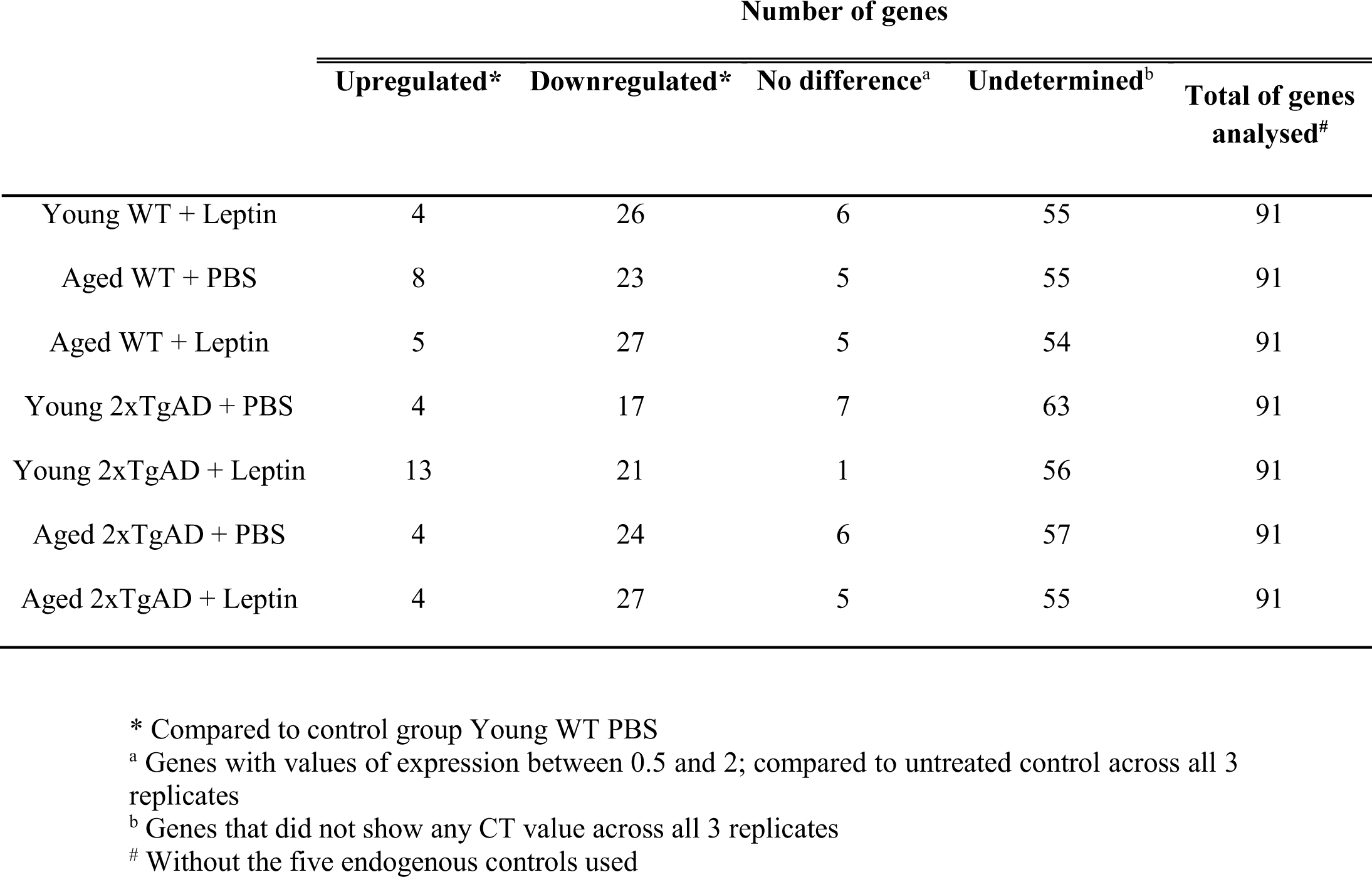
Number of genes related to oxidative stress in the hippocampus that were upregulated by leptin above a 2.0 ratio or were downregulated by leptin below a 0.5 ratio.

**TABLE 2:**
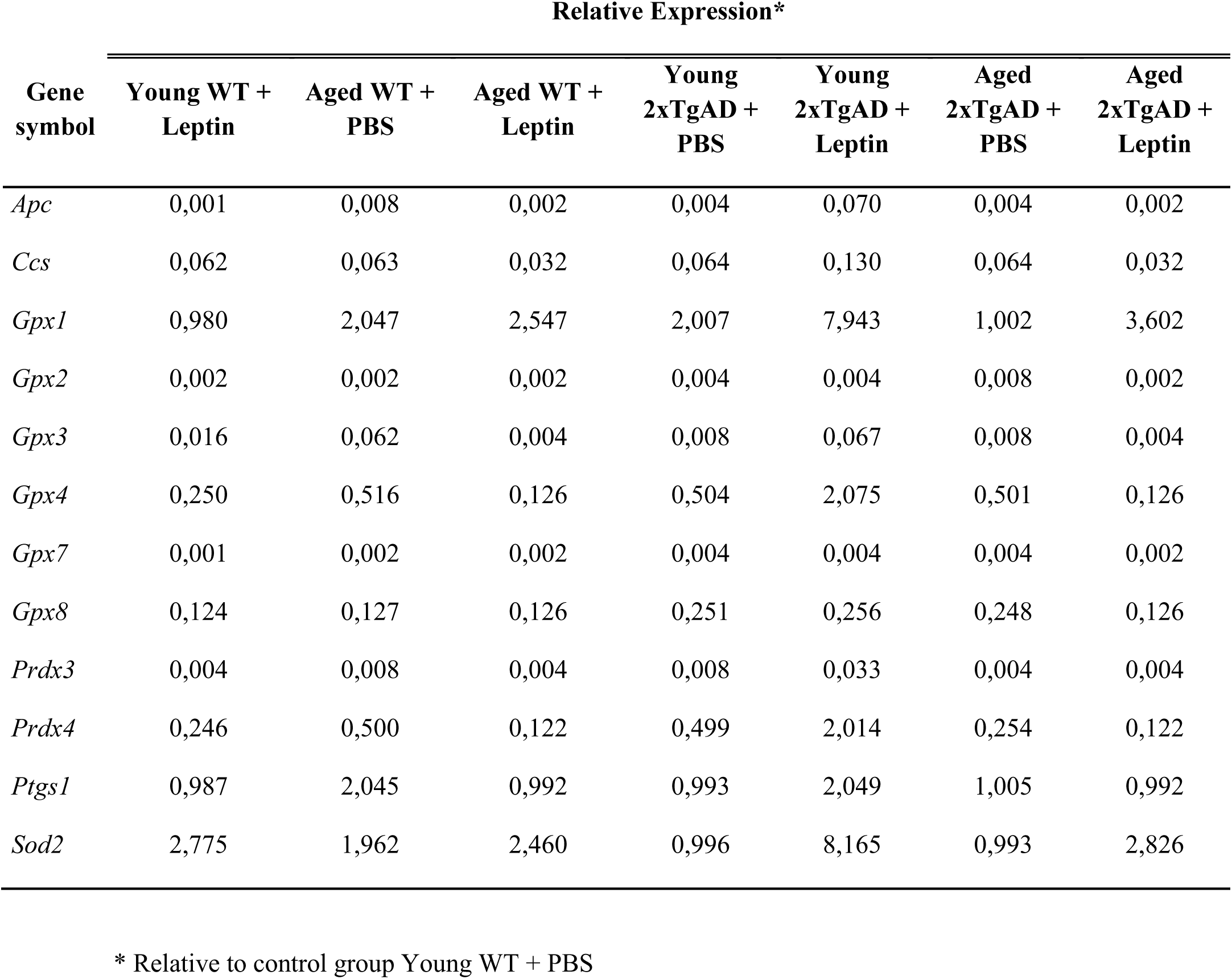
Fold increase of the relative expression (RE) of genes related to antioxidant enzymes that were upregulated by leptin

**Figure 6.**
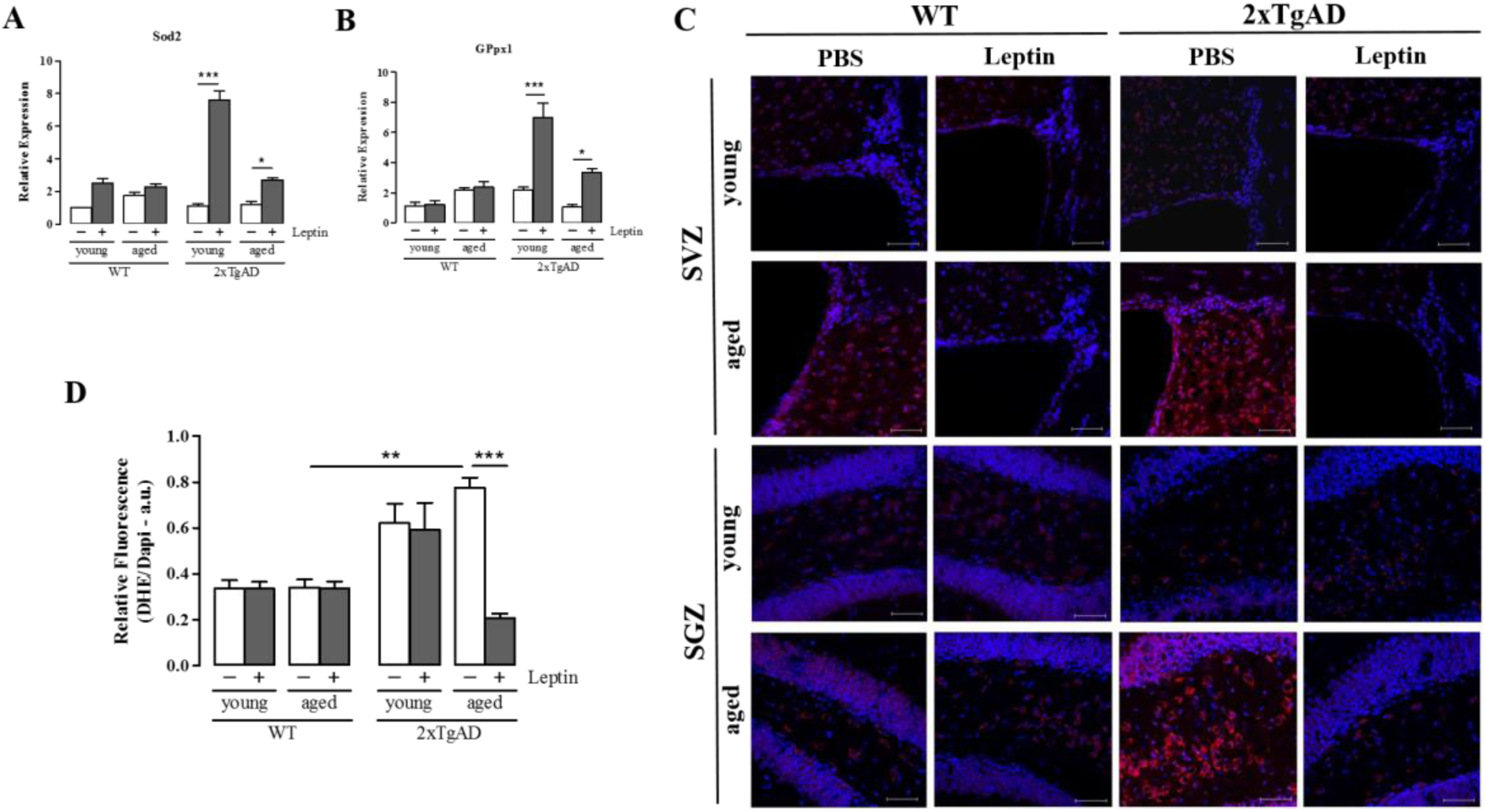
Leptin increases Sod2 and GPx1 and reduces oxidative stress in the hippocampus of 2xTgAD mice. In vivo leptin administration for 7 days upregulated the expression of Sod2 **(A)** and GPx1 gene expression **(B)** in the hippocampus of 2xTgAD but not in wild type. The data were normalized using four endogenous controls and are relative to young WT PBS control group. Data are expressed as mean ± SEM, *n* = 5 animals per group. \**p* < 0.05; \*\**p* < 0.01. **(C)** Representative confocal images of superoxide staining in the hippocampus of WT and 2xTgAD animals treated and not treated with leptin for 7 days. Dihydroethidium staining (DHE - red). The nuclei were stained with DAPI (blue). The ratio intensity of DHE / DAPI fluorescence was used to calculate the oxidative stress rate. Intense levels of DHE indicate greater occurrence of superoxide anion formation. **(D)** Quantification of superoxide anion by the analysis of pixilation. A decrease in leptin-treated aged 2xTgAD animals was observed when compared to untreated animals. Scale bar = 20μm. Data are expressed as mean ± SEM represented by error bars, *n* = 6 animals per group. \*\*\**p* < 0.001.

Since our results showed significant differences in *Sod2* expression in leptin-treated AD animals, we focused on analyzing whether the antioxidant activity of leptin could decrease the level of superoxide produced and attenuate the oxidative stress in AD animals. For that, we used dihydroethidium (DHE), which is a detector of intracellular superoxide. Leptin treatment significantly reduced cellular superoxide levels in the hippocampus (Figure 6C, D) of 2xTgAD animals. The indicator of the presence of superoxide production were four times greater in the dentate gyrus of untreated aged 2xTgAD than in treated animals, that exhibited a significant decrease in the DHE/DAPI ratio, confirming the protective role of this hormone. Curiously in SVZ, leptin decreased superoxide production also in younger animals.

Taken together, these results suggest that *Sod2* protects the aging brain against oxidative stress in AD by controlling superoxide overproduction and that leptin treatment can restore this antioxidant pattern.

### Leptin reduces Aβ plaque accumulation and neurodegeneration

Leptin has been shown to reduce brain Aβ levels in 6-months old CRND8 transgenic mice following 8 weeks of treatment (Greco et al., 2010). Our data show that Aβ staining in the hippocampus of aged 2xTgAD mice was significantly reduced after treatment with leptin (Figure 7B).

**Figure 7.**
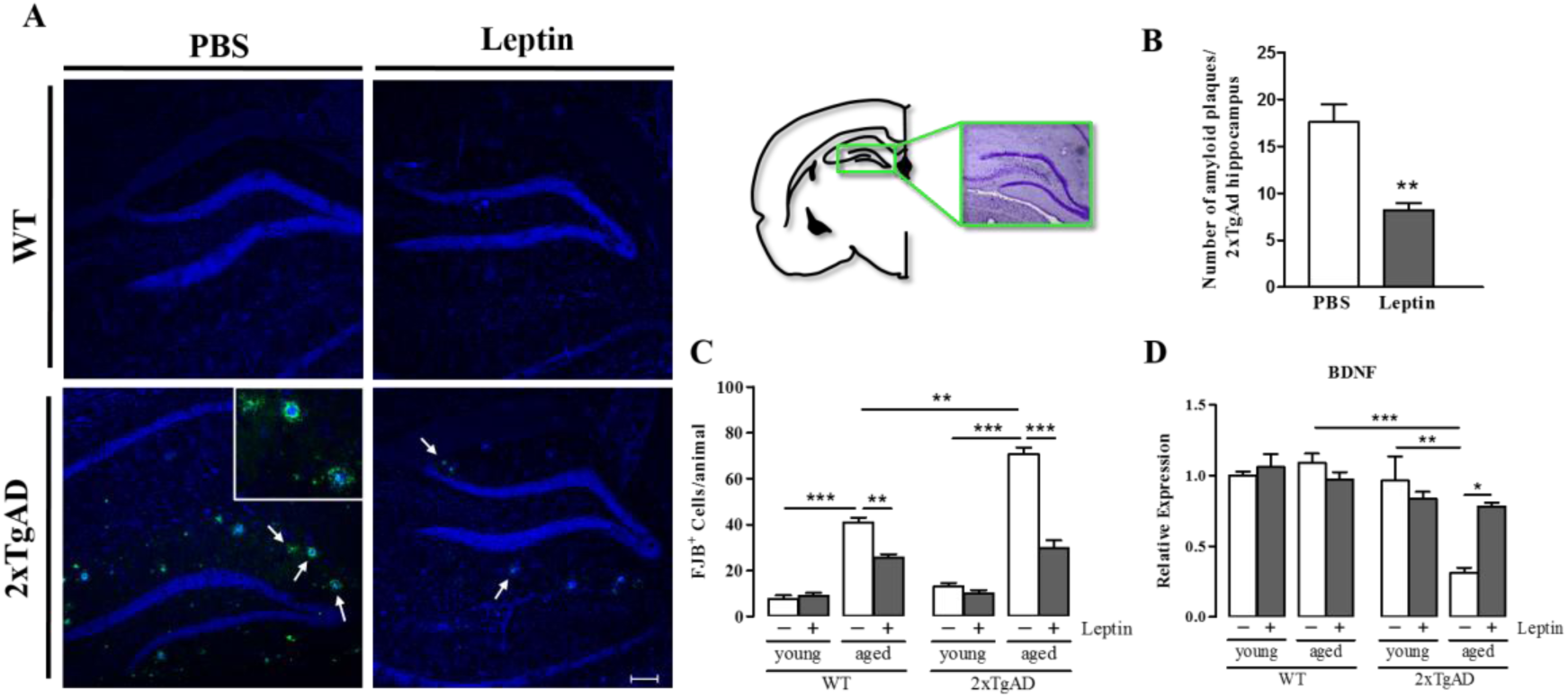
Leptin decreases accumulation of Aβ in the hippocampus and increases BDNF expression. Leptin-treated 2xTgAD mice exhibit reduced levels of Aβ deposits (green) compared with control (PBS) 2xTgAD mice. **(A)** Representative images of Aβ staining in the hippocampus of aged WT and 2xTgAD mice treated with PBS or leptin. Scale bar = 100µm. **(B)** Quantitative analysis of Aβ shows lower number of plaques in the brain of 2xTgAd treated mice. The arrows point at the senile plaques, which are magnified in the inset. Data are expressed as mean ± SEM, *n* = 5 animals per group. \*\**p* < 0.01. **(C)** Quantitative analysis of FJB staining revealed significant reduction in the number of labelled cells after leptin treatment, indicating less neurodegeneration. Aged 2xTgAD animals show downregulation of BDNF when compared with WT animals, and leptin administration restores the levels of BDNF. One-way ANOVA, data are expressed as mean ± SEM, *n* = 6 animals per group. \**p* < 0.05; \*\**p* < 0.01; \*\*\**p* < 0.001.

Taking into account age and lineage of animals, we observed that the number of FJB^+^ cells increases as a function of time and disease in the hippocampus. An example of the contrasting findings between control and leptin groups can be visualized in Figure 7C. In fact, we observed that Fluoro-Jade B (FJB) labeling in the hippocampus of 2xTgAD mice decreased significantly after treatment with leptin as compared with control group and was similar to WT mice (See Supplental Data Figure S2). In addition, the hippocampus of aged transgenic mice contains more neurodegenerating cells when compared to young mice and to transgenic mice treated with leptin. We also observed FJB^+^ staining in other areas, such as cortical regions and thalamus (data not shown). Due to the association between brain-derived neurotrophic factor (BDNF) levels and the onset of AD, we investigated BDNF mRNA expression in the animals. BDNF was detected in all groups, with significant increments in aged 2xTgAD leptin-treated group while lower levels of BDNF expression were associated with the progression of pathology (Figure 7D).

These results suggest a novel mechanism linking leptin action to BDNF expression in AD, while it also implies the neurogenesis in this pathology.

### Leptin treatment attenuates microglia activation in the hippocampus

To verify neuroinflammation in the hippocampus, microglia was immunolabeled with anti-Iba1 and counted (Figure 9A). Leptin treatment significantly reduced microglia burden in 2xTgAD mice (Figure 9B). After 7 days of treatment with leptin, a few randomly distributed Iba1^+^ cells were observed in WT animals while in 2xTgAD treated mice, the remaining Iba1^+^ cells were observed mainly at sites of amyloid-beta plaques (Figure 8 and 9C). Beside this, the ramified phenotype of microglia predominated in the great majority of 2xTgAD animals, showing that the cytoplasmic processes of Iba1^+^ cells can serve as a sensor to explore the surrounding environment, including the of beta-amyloid plaques.

**Figure 8.**
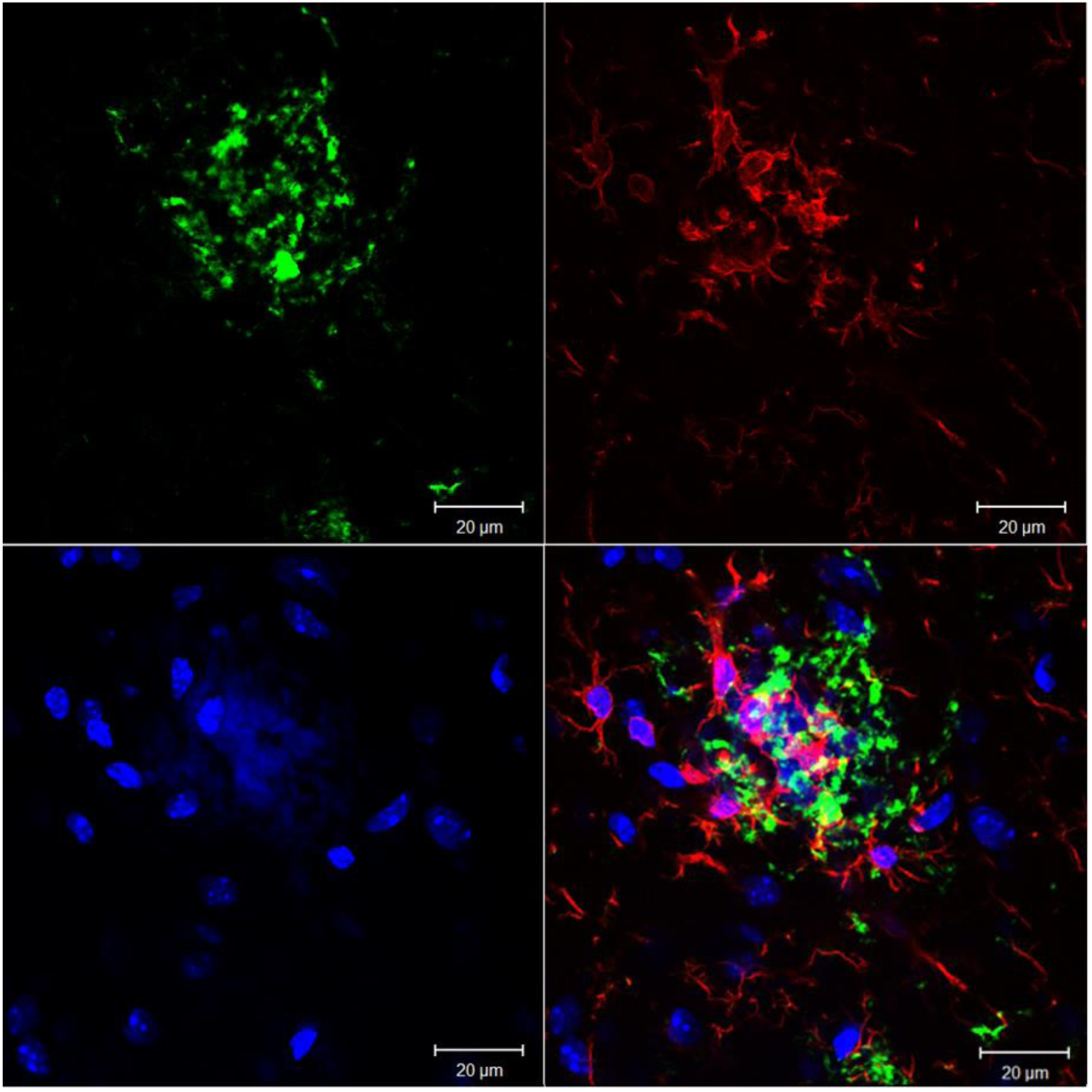
Activated microglia on amyloid plaque. Scale bar = 20 µm.

**Figure 9.**
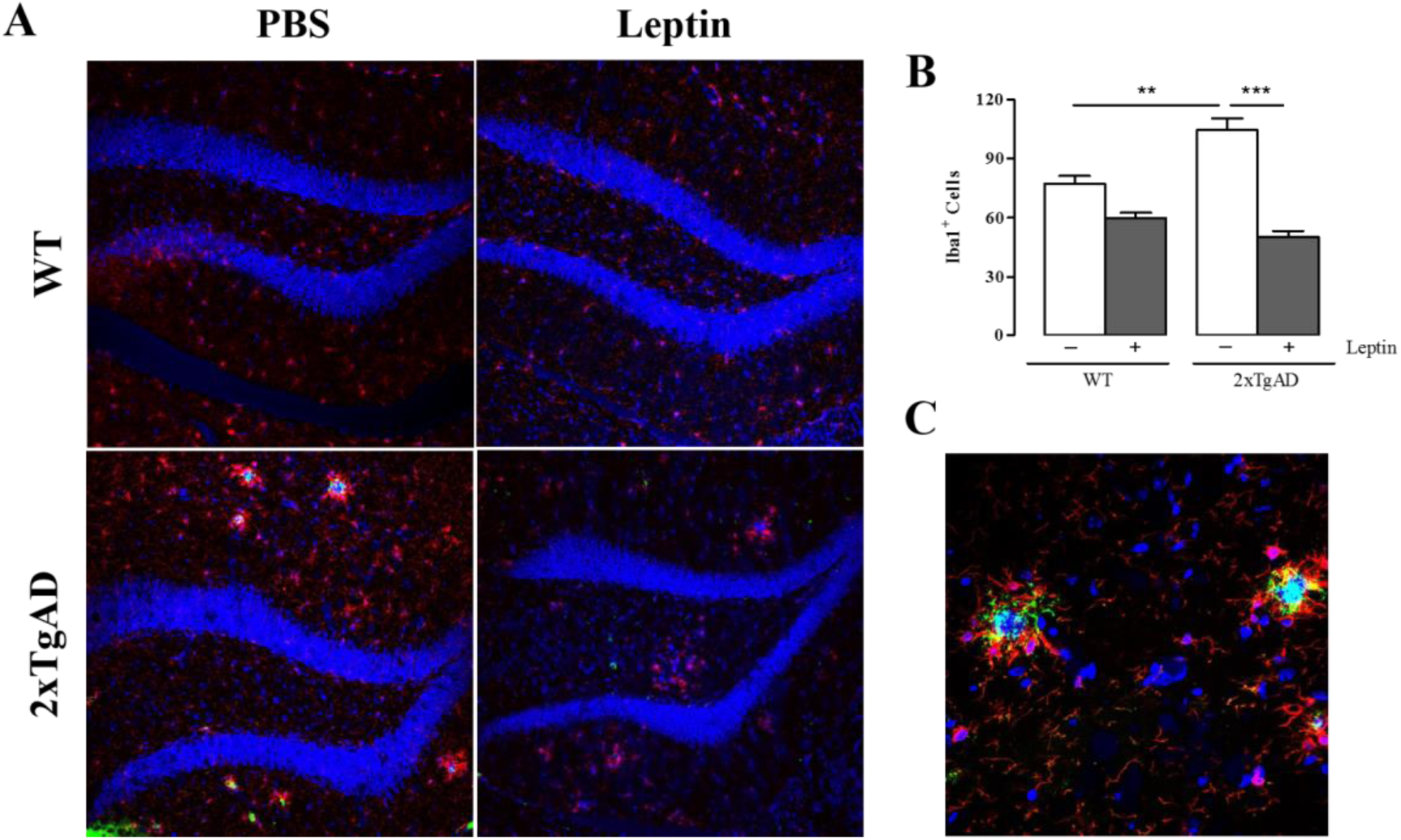
Leptin attenuate microglia activation in the hippocampus of WT and 2xTgAD mice. **(A)** Representative images of Iba-1^+^ cells, in the hippocampus sections of WT and 2xTgAD animals) yellow arrow indicates the morphology of activated microglia **(B)** Quantitative analysis of Iba-1^+^ stained cells. **(C**) Higher magnification of amyloid plaques seen in 2xTgAD - PBS animal. One-way ANOVA, data are expressed as mean ± SEM, *n* = 6 mice per group, 4-8 sections per animal; * p < 0.05; *** p < 0.001. Scale bar = 100μm in (A) and 20μm in (C).

## DISCUSSION

Leptin is known to suppress appetite and reduce body weight gain (Zhang et al., 1994) and have been shown to enhance hippocampal neurogenesis (Garza et al., 2008; Pérez-González et al., 2011). Since dietary restriction and physical activity have been shown to improve hippocampal neurogenesis, one possibility is that the effects of leptin on neurogenesis could be mediated by its action in food intake and metabolism regulation (Lee et al., 2002; Trejo et al., 2001). However, in our study, the treatment with leptin had no effect on the weight of treated animals. Garza et al. (2008) reported that body weight gain decreased without significant effect on hippocampal neurogenesis after acute leptin injection. Moreover, it has been demonstrated that locomotor activity was not altered by leptin chronic administration (Overton et al., 2001). These findings support that the effect of leptin on neurogenesis is dissociated from its impact on energy homeostasis.

In the present study, we have demonstrated that administration of leptin to 2xTgAD mice increases proliferation of neural progenitors in the main neurogenic niches and reduces neurodegeneration, thereby suggesting by this mechanism that dead or damaged neurons could be replaced.

Prior studies have shown that neurogenesis occurs in the adult mammalian brain, although at a reduced rate with advancing of age, despite the fact that aged brains partially retain neurogenesis capacity through different mechanisms (Kempermann et al., 2002). In conditions like AD, which occurs at an increasing frequency with advancing age, the ability of the aged brain to mobilize new neurons opens new possibilities for cell**-** replacement therapy. However, an increase in neurogenesis may not be enough to compensate the progressive pathological changes that occur in this disease (Jin et al., 2004). Neurogenesis is a process that can be stimulated by physiological factors, such as growth factors and environmental enrichment, and by pathological processes, including neurodegeneration and trauma (Pérez-González et al., 2011). *In vitro* and *in vivo* studies have shown that leptin stimulates hippocampal neurogenesis by increasing cell proliferation (Garza et al., 2008) and neuroprotection (Weng et al., 2007; Zhang et al., 2007).

It is well known that the microenvironment of the AD brain is toxic to neurons due to the formation of Aβ oligomers (Rapoport et al., 2002). This may be one of the reasons why there is a limitation in the ability to repair neuronal death through neurogenesis in AD. Our findings presented here indicate that leptin administration reduces the accumulation of Aβ deposits in 12 months-age 2xTgAd mice, coinciding with previous published studies (Pérez-González et al, 2011).

It has been reported that chronic intra-ventricular infusion of leptin protects hippocampal neurons from cell death induced by neuronal insults (Guo et al., 2008). In this sense, our data are consistent with the findings of Pérez-González et al. (2011) who used lentivirus gene delivery to overexpress leptin in the brains of AD animals. Interestingly they also observed that leptin significantly reduced neuronal degeneration in the hippocampus of AD model, also suggesting that leptin has a neuroprotective effect.

In the present study, we also evaluated the action of leptin on gene expression of antioxidant enzymes. In addition, we checked whether leptin was able to decrease previously existing levels of superoxide in the neurogenic niches of AD animals. Although the main cause of neuronal death in AD is the formation of senile plaques, *in vivo* evidence showed that mitochondrial dysfunction and redox imbalance triggers or worsens AD-related neuronal deficits (Calon et al.,2004; Lustbader et al., 2004; Beal, 2005). Studies suggest that there are multiple mechanisms by which oxidative stress may accumulate and create dysfunctional neuronal responses in AD and that development of the AD phenotype requires multiple insults (Zhu et al., 2004). Since our previous results demonstrate the ability of leptin to decrease neurodegeneration in animal models of AD, we decided to investigate whether leptin could improve the existing oxidative stress profile in this pathology. We have demonstrated the existence of significantly abnormal increased superoxide generation and the downregulation of *Sod2* expression in hippocampus of aged 2xTgAD, suggesting a negative correlation between cellular superoxide production and *Sod2* decline in advanced AD. We also found that the fluorescence intensity of DHE staining was increased in non-treated aged 2xTgAD in comparison to WT and that the superoxide levels decreased after treatment with leptin in these animals, in both niches analyzed. Besides, no difference was found in WT group. Surprisingly, in SVZ, the antioxidant action of leptin was detected in younger animals, whereas in the hippocampus, the same result could only be observed in the aged animals.

We studied gene expression of some proliferation and neural markers expressed in brain asked whether they were modified in the transgenic mice and if leptin could alter this pattern. Leptin induced increased expression of DCX, a highly expressed gene in neuroblasts, which was found at low levels in all groups of 2xTgAD animals prior to treatment. Increased PCNA expression was only observed in WT animals of both ages, whereas increased Ki67 and nestin expression was also be observed in transgenic animals. Unlike expected by the effect of leptin on cell proliferation, this hormone decreased the gene expression of GFAP, which was twice as high in the aged 2xTgAD animals compared to WT and treated animals. This fact suggests that the decrease in GFAP is linked to the reduction of astrogliosis observed in aged AD animals. No differences were observed in the expression of genes makers of mature neurons (βIII-tubulin, Map2a and NeuN) when leptin was used, and lower expression was observed in the aged 2xTgAD animals compared to WT mice.

Furthermore, our data show that leptin enhanced adult neurogenesis both in SVZ and SGZ of WT animals and models of AD. The leptin-stimulated neurogenesis resulted not only in increased proliferation of neural progenitors in all analyzed groups but also in the decrease of astrogliosis observed in aged 2xTgAD animals.

In order to determine whether leptin could also affect differentiation of neural stem cells in neuroblasts or astrocytes, or even affect proliferation of neural progenitors, cell differentiation was assessed using DCX and GFAP labeling. Our results also showed increased expression of the neuronal marker DCX in the SVZ and hippocampus of mice. With the use of immunohistochemistry, DCX was shown to localize in the SGZ, which is associated with neurogenesis, and involved in the pathogenesis of AD.

Chronic neuroinflammation is common to nearly all neurodegenerative diseases, and it contributes to their pathophysiology (Heneka et al., 2014). Astrocyte cells are capable of local proliferation under certain conditions, with indications that their progeny may resume normal physiological function over time (Bardehle et al., 2013; Buffo et al., 2008). It is known that injury and degeneration can cause astrogliosis and subsequent astrocyte proliferation, although the exact molecular stimulus of proliferation is currently unknown (Doetsch, 2003). Nevertheless, although anti-inflammatory and immunosuppressive therapies have demonstrated some efficacy in neurodegenerative disease models, these treatments have largely failed in the clinic (Arvanitakis et al., 2008; Wyss-Coray et al., 2012). Our data show that leptin treatment may decrease reactive astrogliosis in aged 2xTgAD animals as seen by the exacerbated proliferation in the hippocampus of these animals which is subsequent to the neurodegenerative lesions found in this pathology. Our findings are consistent with the results of Garcia et al. (2014), which observed a large number of GFAP^+^ cells in the hippocampus of this animal model at the age of 9 and 12 months, compared to WT mice.

In the present study, we observed neuroinflammation in the hippocampal area, including microgliosis and astrogliosis. A recent study mentions that Iba1 is widely expressed in microglia of the hippocampus and cortex of 12-month-old 2xTgAD animals (Unger eta l., 2018). In AD mice, these cells clustered primarily at sites of amyloid plaques (Nimmerjahn et al., 2005; Unger eta l., 2018) supporting our interpretation of Iba-1 stainning in the hippocampus of 2xTgAD mice (Figure 8). We observed that leptin can attenuate Iba-1 expression, compared with 2xTgAD saline injected group (Figure 9B). On the other hand, in the 2xTgAD PBS group, Iba-1 immunoreactive microglia were similar to those in the WT group. This corroborates a previous publication that attests that leptin decreases the level of activated microglia in the hippocampus 1 day after transient ischemia (Yan et al., 2011), indicating that this hormone is also effective in reducing this neuroinflammatory marker. Promisingly, microglia activation seems to be an important target reached by leptin treatement.

Therapeutic explorations on AD have been constantly conducted on the pathogenesis and subsequently tailored therapeutic intervention, especially in neurobiological field, and the discovery of neurotrophins (NTs) was a milestone, which provided a new perspective into neurogenesis and neuron survival (Bothwell, 2014). Brain-derived neurotrophic factor (BDNF) is the most widely distributed NT in adult brain and has key roles in neuronal survival, synaptic plasticity and memory storage in hippocampus (Lu et al., 2013). More recently, it has also been described as important in the central control of energy homeostasis (Rios, 2013; Vanevski and Xu, 2013). BDNF is widely distributed in the brain, being highly expressed in the hippocampus, cortex, and basal forebrain, where it is important for learning, memory, and higher thinking (Huang and Reichardt, 2001; Yamada et al., 2003; Voineskos et al., 2011) and in several hypothalamic nuclei, where it also regulates feeding and body weight (Rios, 2013). Although BDNF is needed in the developmental stages, BDNF levels have been shown to decrease in tissues with aging (Tapia-Aranciba et.al.,2008). In humans, decreased hippocampal volume are associated with decreasing BDNF levels, suggesting there is a relationship that might explain some of the cognitive decline that occurs during aging (Erickson et. al., 2010; Buchman et al., 2016). Changes of BDNF level and mRNA expression have been also reported in brain samples as well as blood of AD patients, which also indicates a potential role of BDNF in the pathogenic process of AD (Peng et al., 2009; Angelucci et al., 2010; Laske et al., 2011; Voineskos et al., 2011; Huan et al., 2012; Francis et al., 2012; Nagata et al., 2013; Ventriglia et al., 2013; Prakash and Kumar, 2014; Song et al., 2015; Li et al., 2017; Marsh and Blurton-Jones, 2017). Correspondingly, higher blood BDNF levels seem to delay the rate of cognitive decline in AD (Bollen et al., 2013). As early as 1991, it was found that BDNF mRNA decreased in the hippocampus of individuals with AD, which suggested that BDNF may be related and contribute to the progression of cell loss in AD (Phillips et al., 1991).

Compared with aged controls, BDNF protein expression as well as mRNA levels were found decreased in hippocampus and frontal, parietal, and temporal cortex of AD postmortem samples (Connor et al., 1997; Hock et al. 2000; Holsinger et al., 2000; Michalski and Fahnestock, 2003; Peng et al., 2005), meanwhile, similar results are also replicated in AD animal models (Peng et al., 2009; Li et al., 2009; Kim et al., 2010; Naert and Rivest, 2012; Maioli et al., 2012; Francis et al., 2012; Meng et al., 2013). In addition, another study demonstrated that induction of Alzheimer’s disease model leads to decrease in BDNF mRNA levels in the hippocampus (Smith et al. 1995).

BDNF functions depend not only on its site of action but also on a complex array of metabolic and environmental factors that regulate its expression in a brain region-specific manner. It is currently accepted that peripheral metabolic cues such as leptin increase BDNF mRNA expression in the nucleus of the hypothalamus. Leptin is an adipocyte-derived hormone that acts as an anorexigenic stimulus and it is known to acutely induce BDNF gene expression in the ventro-medial nucleous of the hypothalamus of fasted mice (Komori et al., 2006; Unger et al., 2007) but is unclear whether leptin regulate mRNA BDNF in the hippocampus, as a modulator of neural activation. In our experiments, changes in hippocampal BDNF gene expression were present in the aged treated group in which significantly higher levels were observed. However, it is unlikely that leptin alone contributed to the increase in BDNF expression observed in this animals.

These results demonstrate the utility of leptin as a potential therapeutic approach for AD, although essential features are needed to ensure its translation to the clinic in the near future.

In summary, our results suggest that leptin increases the production of newborn cells in two neurogenic niches even in aged AD brain. These results support a novel role of leptin in the processes of adult neurogenesis and neurodegeneration, providing new insights into the mechanisms of neurogenic regulation in dementia. Future work will be needed to determine which signaling mechanisms are involved in the neurogenic and neuroprotective function of this hormone. Our data also provide the basis for further analysis of the role of leptin, as an alternative or adjuvant, to slow down or delay the central pathology associated with AD.

## ACKNOWLEDGMENTS

We would like to thank Professor Beatriz Monteiro for 6E10 antibody and Clivandir Severino for technical assistance. This work was supported by Fundação de Amparo à Pesquisa do Estado de São Paulo – FAPESP (2014/24341-9 and 2015/19231-8), Conselho Nacional de Desenvolvimento Científico e Tecnológico – CNPq (404646/2012-3), and Coordenação de Aperfeiçoamento de Pessoal de Nível Superior – CAPES (Finance Code 001).

## AUTHOR CONTRIBUTIONS

Michele L. Calió conceptualized the project, carried out the immunohistochemistry, qPCR experiments, performed all statistical analysis, interpreted the data, and wrote the manuscript. Darci S. Marinho performed DNA extraction and genotyped all the animals, separating them for the experiments. Geisa N. Salles carried out serum extraction, ELISA experiments and tissue preparations. Amanda F. Mosini contributed to RNA and cDNA for molecular analysis and qPCR experiments. Fernando H. Massinhani, performed morphometrical data collection and analysis and assisted cell quantification. Gui M. Ko selected and crossed the heterozygous animals and maintained the colony of transgenic mice and assisted the treatment of animals. Marimélia A. Porcionatto conceptualized the project, designed experiments, interpreted the data, provided guidance and supervision, and wrote the manuscript. The work presented here was carried out in collaboration between all authors. All authors revised the manuscript.

## EXPERIMENTAL PROCEDURES AND DETAILED METHODS

### Animals

Three-month-old (young) and twelve-month-old (aged) male C57BL/6J mice (wild type – WT) and heterozygous double transgenic AβPP/PS1 mice (2xTgAD) - a cross between Tg2576 (overexpressing human AβPP695) and mutant PS1 (M146L) - colony B6.Cg-Tg(APPswe, PSEN1dE9)85Dbo/J (from Jackson Laboratory) - were used. The animals, weighing 25–30g, were purchased from Center of Development of Experimental Models for Biology (CEDEME– UNIFESP), at the *Universidade Federal de Sâo Paulo*, São Paulo, Brazil. Mice were genotyped to confirm the presence of the transgenes. Animals were housed in groups of five and maintained on a 12-h light/dark cycle (lights on at 06:00 h) with *ad libitum* access to food and water. Animals were habituated to the housing condition for 7–10 days prior to the beginning of the experimental procedures. All procedures were carried out in accordance with the internal Ethical Committee on Animal Research of the Universidade Federal de São Paulo UNIFESP/EPM (CEP 6882070814).

### Drugs

Recombinant mouse leptin (R&D Systems, Minneapolis, MN) was dissolved in sterile saline solution.

### Leptin treatment

Leptin at a dose of 1mg/kg was used to study its effect on cell proliferation in the neurogenic niches. This dose of leptin was chosen based upon its effectiveness in inducing neurogenesis in mice (Garza et al., 2008). Animals of both ages and lineages received intraperitoneal (i.p.) injection of saline (PBS) or leptin (1mg/kg), once a day at the end of the light cycle for 7 consecutive days (n = 6 animals per group). Body weight was measured daily during the period of treatment. In the last two days of treatment, mice were injected with bromodeoxyuridine (BrdU), a thymidine analog that can be incorporated into the DNA during the S-phase of cell division, to label proliferating cells and determine the fate of newly proliferated cells. Animals received four intraperitoneal injections of BrdU in total (50mg/kg body weight in 0.1M PBS, pH7.2), spacing at 8-h intervals over 48 h. The time interval corresponds with the length of S-phase of cell division reported in mice. Animals were transcardially perfused 8h after the last BrdU injection for immunohistochemical evaluation of BrdU-labeled cells.

### Biochemical measurements

Serum samples were obtained from aged animals for correlating the decrease in plasma leptin levels and the onset of AD and stored at –80°C. The concentrations of leptin levels from non-transgenic WT and 2xTgAD mice were determined using the commercial ELISA kit Quantikine Mouse Leptin Immunoassay (R&D Systems; Minneapolis, MN, USA) according to the manufacturer’s recommendations.

### Immunohistochemistry Tissue Preparation

Animals were anesthetized with an intramuscular injection of ketamine cocktail (43mg/kg ketamine, 10mg/kg xylazine, and 1.4mg/kg acepromazine in saline) and perfused through the ascending aorta using 0.1M PBS followed by 4% paraformaldehyde (PFA) in PBS. The brain was removed from skull and fixed overnight in 4% PFA, before being transferred to 30% sucrose solution for 48h.. Brains were embedded in frozen tisuue embedding media(HistoPrep TM, Fisher Scientific) and coronally crypsectioned at 30µm sections and stored in cryoprotectant (30% sucrose, 30% ethylene glycol, 0.05M sodium phosphate buffer) until processing for immunohistochemistry.

### BrdU Immunostaining

Once BrdU is incorporated into the DNA, the sections were first pretreated to denature DNA to make it assessable to BrdU antibody. Brain sections were briefly rinsed in PBS buffer, treated with 1% hydrogen peroxide for 10 min, and incubated in 2N HCl solution with 0.3% Triton X-100 for 30 min followed by incubation with 0.1M boric acid for 10 min. Subsequently, the sections were blocked in a solution containing 0.3% Triton X-100 and 3% normal goat serum in PBS for 1h, and then incubated with rat monoclonal anti-BrdU (anti-Rat, 1:300, Molecular Probes) in the blocking solution overnight at 4°C. After rinsing in PBS buffer, the sections were then incubated with goat anti-rat Alexa Fluor 647nm (1:300, Molecular Probes) in blocking solution for 1h at room temperature. A negative sample test was performed using serum rather than primary antibody to establish the specificity of the immunostaining. The slides were examined under a laser scanning confocal microscope (Zeiss LSM 510 Axiovert; Carl Zeiss GmbH, Germany).

### Fluoro-Jade B labeling

FJB is a fluorochrome derived from fluorescein, and is commonly used in neuroscience to label degenerating neurons in brain tissue. Fluoro-Jade B (Histochem, Jefferson, AR) staining was carried out as described by Schmued and Hopkins (2000). Briefly, paraformaldehyde-fixed brain sections were mounted on 1.5% gelatin-coated slides, air-dried overnight at room temperature and then for 30 min at 40°C before staining. Sections were immersed for 5 min in a solution containing 1% sodium hydroxide in 80% alcohol, then for 2 min in 70% ethanol, and finally for 1 min in distilled water. Sections were then oxidized by immersion for 10 min in 0.06% KMnO4, under moderate shaking. After several rinses in distilled water, sections were incubated for 30 min in 0.004% FJB dye in 0.1% acetic acid, rinsed thoroughly in distilled water, and placed into a heater set to 40°C until the tissue was completely dry. Finally, they were cleared in xylene and coverslipped using D.P.X. mounting medium (Sigma).

### Fluorescence Immunolabeling

To determine the phenotypes of BrdU-labeled cells and amyloid-β deposits, brain sections were processed for doublecortin (DCX), glial fibrillary acidic protein (GFAP), Leptin-receptor (LepRb), Aβ (amyloid beta) or Iba1 (ionized calcium-binding adaptor molecule) fluorescent double or triple labeling. Briefly, free-floating SVZ and hippocampus sections were collected and first pretreated for DNA denaturation as described above, and then incubated with rat-anti-BrdU antibody (1:300, Molecular Probes) and guinea pig anti-DCX (1:500, Merk Milipore), chicken anti-GFAP (1:500, Merk, Milipore) or rabbit anti-LepRb (AbCam). To detect Aβ deposits, brain sections from mice were immunostained without DNA denaturation step and the following primary antibody were used: mouse anti-β-Amyloid 6E10 (1:1000, Covance/Biologend). After washing in PBS (0.1 M) buffer three times, sections were incubated for 1h with fluorescent secondary antibodies conjugated with Alexa Fluor 647 goat anti-rat IgG to reveal immunoreactivity of BrdU, Alexa Fluor 488 goat anti-guinea pig IgG to reveal DCX or Alexa Fluor 594 goat anti-chicken IgG to reveal immunoreactivity of GFAP or LepR, respectively (1:300 for all three antibodies, Molecular Probes, Eugene, Ontario, Canada) and Alexa Fluor 488 goat anti-mouse IgG to reveal Aβ plaques. Then, the sections were rinsed in PBS and mounted onto coated glass slides, and coversliped with fluorescent mounting medium.

### Superoxide anion detection

Superoxide was detected with the oxidative fluorescent probe DHE (dihydroethidium; Molecular Probes, CA, USA), which reacts with O_2_., resulting ethidine fluorescence. To remove the differences in cellular densities, DAPI was used in all of the frozen slices. The frozen brain sections were incubated in a light-protected humidified chamber at room temperature with DHE (5 mM) for 5 min and then counterstained with the nuclear tracer DAPI (5 mM; Sigma). The slides were immediately analyzed with a computer-based digitizing image system (Zeiss Axiovert 100M; Carl Zeiss, Germany) connected to an LSM 510 Confocal Laser Scanning System. Fluorescence was detected with 510–560 nm excitation and 590 nm emission filters. The results are expressed as the DHE/DAPI ratio according Oudot et al (2006).

### Cell Quantification and Analyses

The SVZ and hippocampus coronal tissue sections were mounted on glass slides and were numerically coded, and stereological counts were performed by experimenters blinded to the experimental condition of each sample. The brain was sectioned at 30µm, and every eight section was spaced 240µm apart throughout the entire SVZ and hippocampus extent was used to assess the number of labeled cells. This strategy makes certain that the same cell will not be counted twice on adjacent sections and guarantees that the area counted for each animal is constant. Counting was accomplished at 40x magnification. This allowed clear visualization of each cell and enabled us to distinguish single cells from clusters. In the SVZ, the cells of the anterolateral region were considered while in the hippocampus cells of the SGZ and the gyrus were counted. Cells were counted throughout each section by focusing through each focal plane of the section to ensure that all labeled cells were visible. The extent of colocalization was validated by viewing cells on three planes (X, Y, and Z) using Z-plane sectioning. Cells single labeled for BrdU and FJB or double labeled for BrdU/DCX, BrdU/GFAP or BrdU/LepR were counted. The percentage of BrdU cells double labeled was calculated by dividing the number of double labeled cells by the total number of BrdU cells and multiplying 100. Morphometrical analysis was performed using ImageJ software (NIH Image).

### Real time quantitative PCR

To determine the action of leptin on gene expression, quantitative reverse transcription PCR (qPCR) was performed. The hippocampi of animals of each group were collected, and total RNA was extracted. The extracted RNA was subsequently used to generate cDNA using SuperScriptTM First Strand Synthesis System for RT-PCR. The cDNA samples were amplified using a 7500 Real-Time PCR System (Applied Biosystems, CA, USA) and TaqMan PCR Master Mix (Applied Biosystems, CA, USA) according to the manufacturer’s instructions. After amplification, the PCR products were analyzed with the ABI PRISM sequence detection software (Applied Biosystems, CA, USA). In addition, melting curve analysis was performed to confirm the authenticity of the PCR products. Quantitative gene expression were analyzed by comparative C(T) method (2^−ΔΔ^CT) according Livak & Schmittgen (2008). TaqMan probes to caspase-3, NeuN, Map2a, DCX, GFAP, βIII Tubulin, Ki67, PCNA, Nestin or Custom TaqMan Array Fast Plates to Oxidative stress were used (all probes and mixes were purchased from Applied Biosystems, CA, USA). The expression of each gene was normalized by the mean of the four endogenous controls (β-actin, 18S, Gapdh and Hprt).

### Statistical Analysis

Comparison of results was performed by one-way analysis of variance (ANOVA) with repeated measures followed by Tukey’s *post hoc* using GraphPad Prism software version 5.0 (GraphPad Software, La Jolla, CA, USA). The data are expressed as mean ± Standard error of the mean (SEM). Significance was set at \**p* < 0.05, \*\**p* < 0.01, \*\*\**p* < 0.001.

## GRAPHICAL ABSTRACT

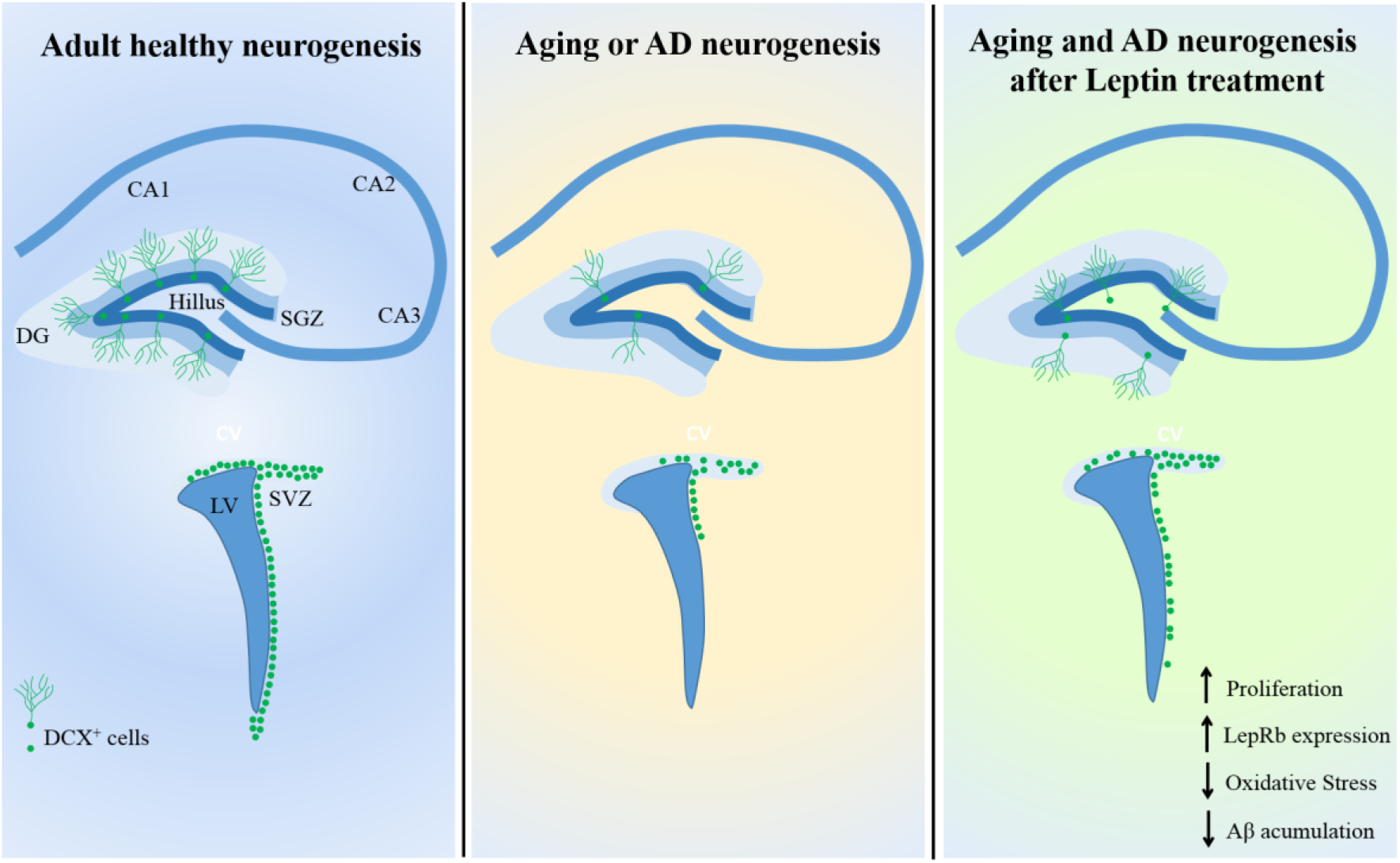

### SUPPLEMENTAL MATERIAL

**TABLE S1:**
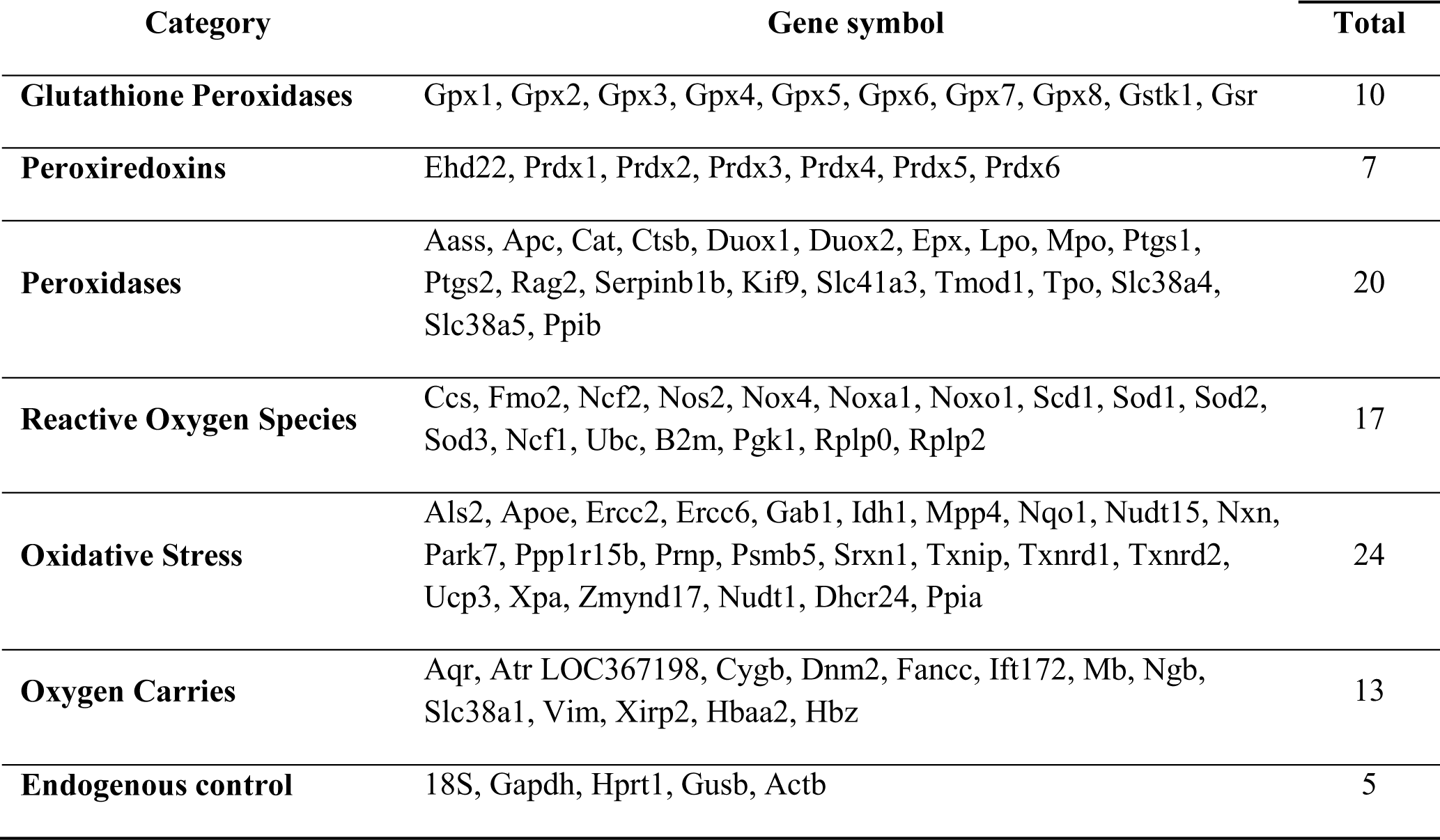
Target genes of Oxidative Stress and Antoxidant defense investigated in hipoccampi of WT and 2xTgAD treated with leptin

**Figure S1.**
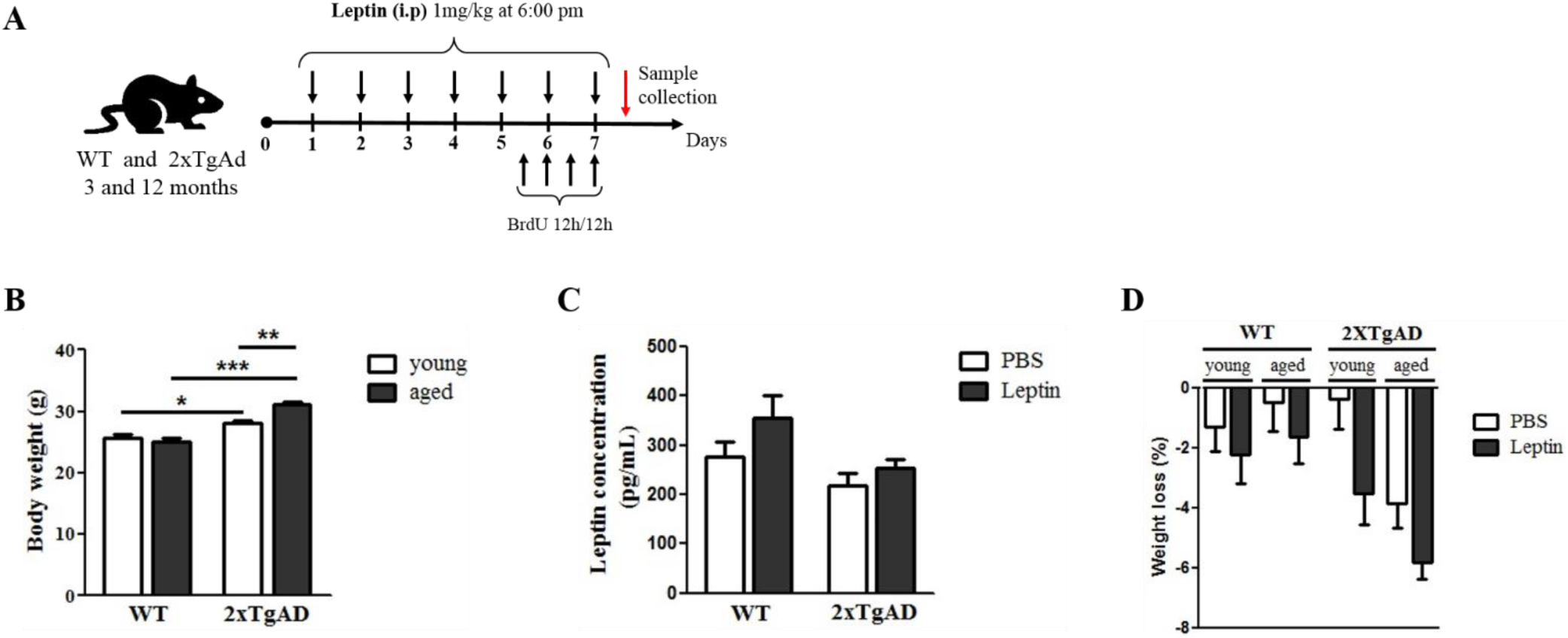
Experimental procedure, body weight and leptin concentration in the sera of aged mice and weight loss. (A) Experimental timeline. (B) Body weight of animals before starting the treatment (*n* 10 per group). Aged 2xTgAD animals show higher body weight when compared to WT and adults 2xTgAD animals. (C) Leptin levels showed a trend for a negative correlation with AD, although this was not statistically significant (*p* = 0.065). (D) Effect of leptin administration on body weight loss. Body weights was measured for 07 days during the treatment and the percentage loss did not show significant variations after the treatment (*n* 5 per group). Data are expressed as mean ± SEM. \*\*\**p* < 0.001; \*\**p* < 0.01; \**p* < 0.05.

**Figure S2.**
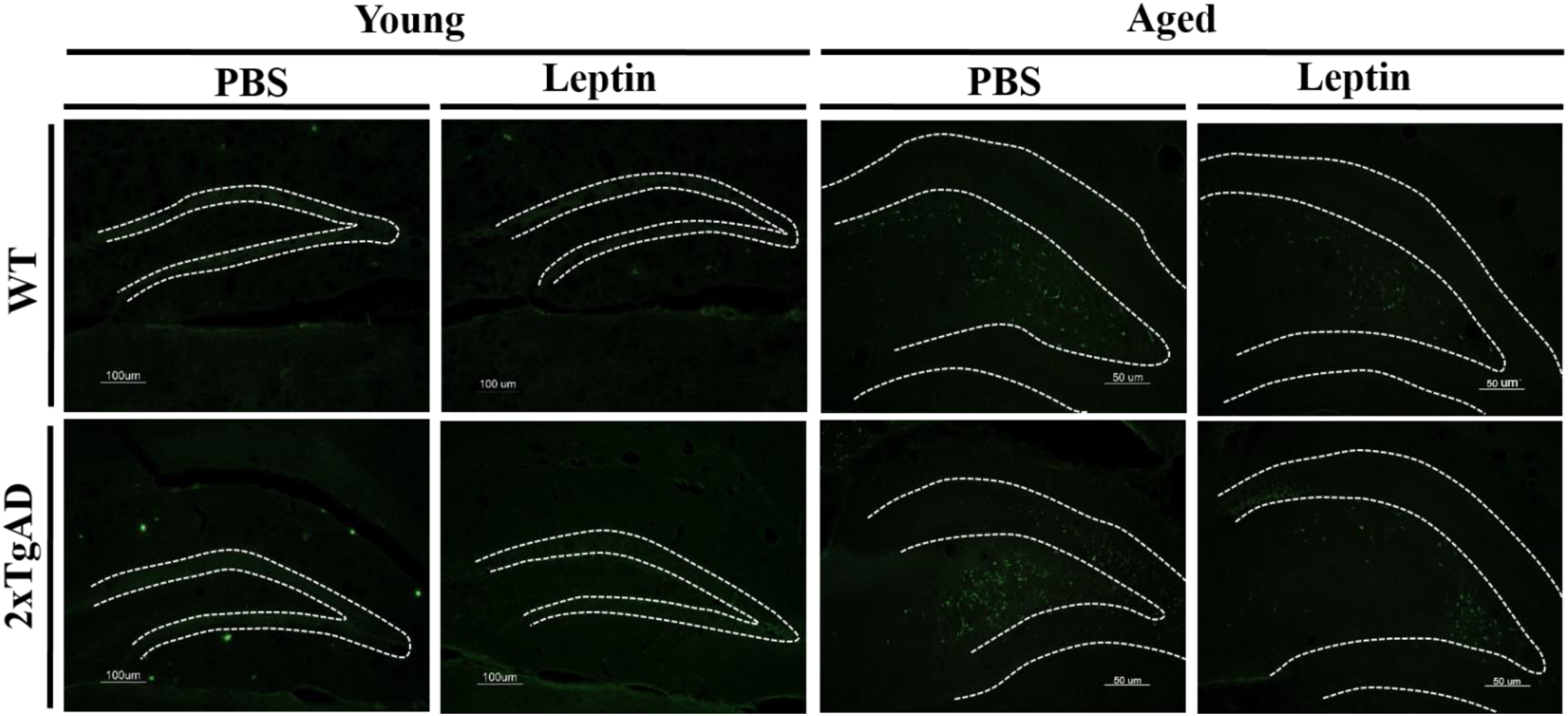
Effect of Leptin on Fluoro-Jade B staining in the hippocampus. Photomicrographs show fluorescent Fluoro-Jade B staining neurodegenerative cells in the hippocampus of mice. Scale bar young animals = 100µm; Scale bar aged animals = 50µm.

