## Supplementary material for "Leptin reduces pathology and increases adult neurogenesis in a transgenic mouse model of Alzheimer’s disease": Supplmental

**Table S1:** Target genes of Oxidative Stress and Antoxidant defense investigated in hipoccampi of WT and 2xTgAD treated with leptin

| Category | Gene symbol | Total |
|---|---|---|
| **Glutathione Peroxidases** | Gpx1, Gpx2, Gpx3, Gpx4, Gpx5, Gpx6, Gpx7, Gpx8, Gstk1, Gsr | 10 |
| **Peroxiredoxins** | Ehd22, Prdx1, Prdx2, Prdx3, Prdx4, Prdx5, Prdx6 | 7 |
| **Peroxidases** | Aass, Apc, Cat, Ctsb, Duox1, Duox2, Epx, Lpo, Mpo, Ptgs1, Ptgs2, Rag2, Serpinb1b, Kif9, Slc41a3, Tmod1, Tpo, Slc38a4, Slc38a5, Ppib | 20 |
| **Reactive Oxygen Species** | Ccs, Fmo2, Ncf2, Nos2, Nox4, Noxa1, Noxo1, Scd1, Sod1, Sod2, Sod3, Ncf1, Ubc, B2m, Pgk1, Rplp0, Rplp2 | 17 |
| **Oxidative Stress** | Als2, Apoe, Ercc2, Ercc6, Gab1, Idh1, Mpp4, Nqo1, Nudt15, Nxn, Park7, Ppp1r15b, Prnp, Psmb5, Srxn1, Txnip, Txnrd1, Txnrd2, Ucp3, Xpa, Zmynd17, Nudt1, Dhcr24, Ppia | 24 |
| **Oxygen Carries** | Aqr, Atr LOC367198, Cygb, Dnm2, Fancc, Ift172, Mb, Ngb, Slc38a1, Vim, Xirp2, Hbaa2, Hbz | 13 |
| **Endogenous control** | 18S, Gapdh, Hprt1, Gusb, Actb | 5 |

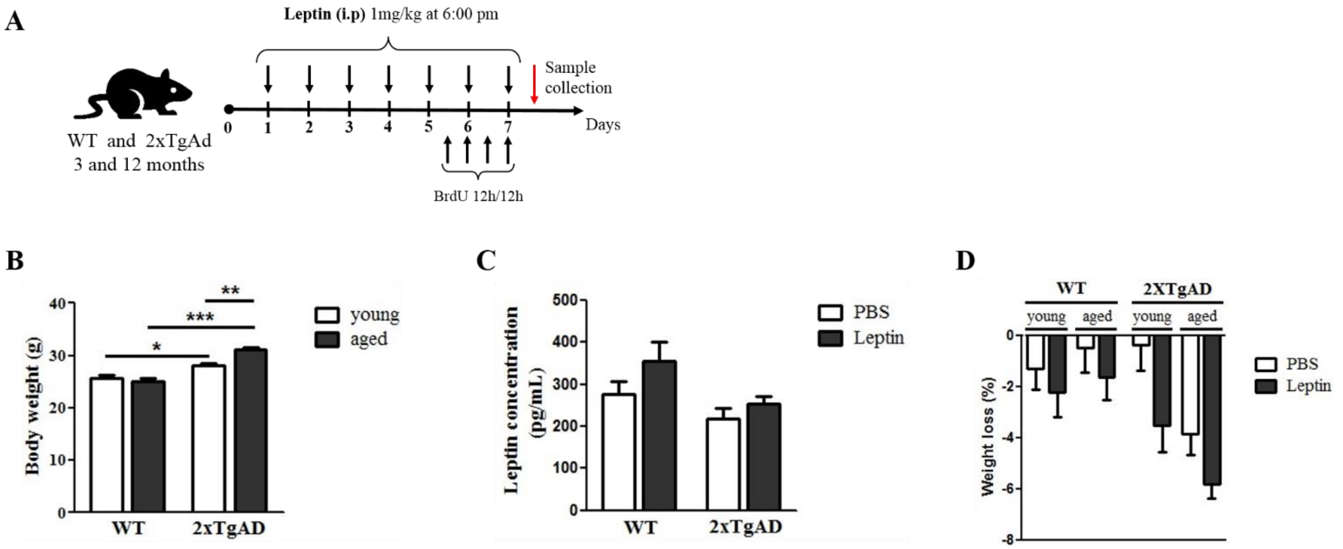


**Figure S1. Experimental procedure, body weight and leptin concentration in the sera of aged mice and weight loss**. (A) Experimental timeline. (B) Body weight of animals before starting the treatment (*n* 10 per group). Aged 2xTgAD animals show higher body weight when compared to WT and adults 2xTgAD animals. (C) Leptin levels showed a trend for a negative correlation with AD, although this was not statistically significant ($p = 0.065$). (D) Effect of leptin administration on body weight loss. Body weights was measured for 07 days during the treatment and the percentage loss did not show significant variations after the treatment (*n* 5 per group). Data are expressed as mean ± SEM. ***$p < 0.001$; **$p < 0.01$; *$p < 0.05$.

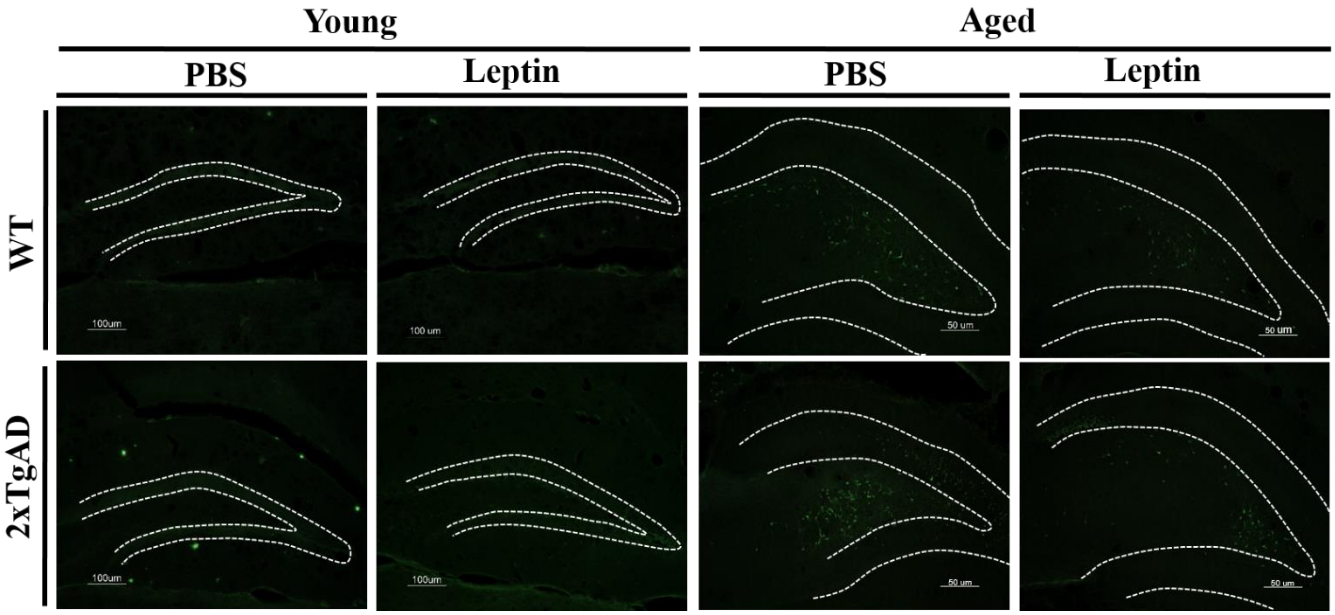


**Figure S2. Effect of Leptin on Fluoro-Jade B staining in the hippocampus.** Photomicrographs show fluorescent Fluoro-Jade B staining neurodegenerative cells in the hippocampus of mice. Scale bar young animals = 100µm; Scale bar aged animals = 50µm.
