## Supplementary material for "Leptin reduces pathology and increases adult neurogenesis in a transgenic mouse model of Alzheimer’s disease": Tables

**Table 1:** Number of genes related to oxidative stress in the hippocampus that were upregulated by leptin above a 2.0 ratio or were downregulated by leptin below a 0.5 ratio.

| | Number of genes | | | | |
|---|---|---|---|---|---|
| | **Upregulated*** | **Downregulated*** | **No difference[a]** | **Undetermined[b]** | **Total of genes analysed[#]** |
| Young WT + Leptin | 4 | 26 | 6 | 55 | 91 |
| Aged WT + PBS | 8 | 23 | 5 | 55 | 91 |
| Aged WT + Leptin | 5 | 27 | 5 | 54 | 91 |
| Young 2xTgAD + PBS | 4 | 17 | 7 | 63 | 91 |
| Young 2xTgAD + Leptin | 13 | 21 | 1 | 56 | 91 |
| Aged 2xTgAD + PBS | 4 | 24 | 6 | 57 | 91 |
| Aged 2xTgAD + Leptin | 4 | 27 | 5 | 55 | 91 |

* Compared to control group Young WT PBS

[a] Genes with values of expression between 0.5 and 2; compared to untreated control across all 3 replicates

[b] Genes that did not show any CT value across all 3 replicates

[#] Without the five endogenous controls used

**Table 2:** Fold increase of the relative expression (RE) of genes related to antioxidant enzymes that were upregulated by leptin

| | Relative Expression* | | | | | | |
|---|---|---|---|---|---|---|---|
| **Gene symbol** | **Young WT + Leptin** | **Aged WT + PBS** | **Aged WT + Leptin** | **Young 2xTgAD + PBS** | **Young 2xTgAD + Leptin** | **Aged 2xTgAD + PBS** | **Aged 2xTgAD + Leptin** |
| *Apc* | 0,001 | 0,008 | 0,002 | 0,004 | 0,070 | 0,004 | 0,002 |
| *Ccs* | 0,062 | 0,063 | 0,032 | 0,064 | 0,130 | 0,064 | 0,032 |
| *Gpx1* | 0,980 | 2,047 | 2,547 | 2,007 | 7,943 | 1,002 | 3,602 |
| *Gpx2* | 0,002 | 0,002 | 0,002 | 0,004 | 0,004 | 0,008 | 0,002 |
| *Gpx3* | 0,016 | 0,062 | 0,004 | 0,008 | 0,067 | 0,008 | 0,004 |
| *Gpx4* | 0,250 | 0,516 | 0,126 | 0,504 | 2,075 | 0,501 | 0,126 |
| *Gpx7* | 0,001 | 0,002 | 0,002 | 0,004 | 0,004 | 0,004 | 0,002 |
| *Gpx8* | 0,124 | 0,127 | 0,126 | 0,251 | 0,256 | 0,248 | 0,126 |
| *Prdx3* | 0,004 | 0,008 | 0,004 | 0,008 | 0,033 | 0,004 | 0,004 |
| *Prdx4* | 0,246 | 0,500 | 0,122 | 0,499 | 2,014 | 0,254 | 0,122 |
| *Ptgs1* | 0,987 | 2,045 | 0,992 | 0,993 | 2,049 | 1,005 | 0,992 |
| *Sod2* | 2,775 | 1,962 | 2,460 | 0,996 | 8,165 | 0,993 | 2,826 |

* Relative to control group Young WT + PBS
